# Mechanisms of blood homeostasis: lineage tracking and a neutral model of cell populations in rhesus macaque

**DOI:** 10.1101/028167

**Authors:** Sidhartha Goyal, Sanggu Kim, Irvin S. Y. Chen, Tom Chou

**Affiliations:** Dept. of Physics, Univ. Toronto, St. George Campus, Toronto, Canada.; Department of Microbiology, Immunology, and Molecular Genetics, UCLA, Los Angeles, USA.; UCLA AIDS Institute and Department of Medicine, UCLA, Los Angeles, USA.; Depts. of Biomathematics and Mathematics, UCLA, Los Angeles, USA.

**Keywords:** hematopoiesis, stem cell clones, lineage tracking, mathematical modeling

## Abstract

How a potentially diverse population of hematopoietic stem cells (HSCs) differentiates and proliferates to supply more than 10^11^ mature blood cells every day in humans remains a key biological question. We investigated this process by quantitatively analyzing the *clonal* structure of peripheral blood that is generated by a population of transplanted lentivirus-marked HSCs in myeloablated rhesus macaques. Each transplanted HSC generates a clonal lineage of cells in the peripheral blood that is then detected and quantified through deep sequencing of the viral vector integration sites (VIS) common within each lineage. This approach allowed us to observe, over a period of 4-12 years, hundreds of distinct clonal lineages. Surprisingly, while the distinct clone sizes varied by three orders of magnitude, we found that collectively, they form a steady-state clone size-distribution with a distinctive shape. Our concise model shows that slow HSC differentiation followed by fast progenitor growth is responsible for the observed broad clone size-distribution. Although all cells are assumed to be statistically identical, analogous to a neutral theory for the different clone lineages, our mathematical approach captures the intrinsic variability in the times to HSC differentiation after transplantation. Steady-state solutions of our model show that the predicted clone size-distribution is sensitive to only two combinations of parameters. By fitting the measured clone size-distributions to our mechanistic model, we estimate both the effective HSC differentiation rate and the number of active HSCs.

## Introduction

Around 10^11^ new mature blood cells are generated in a human every day. Each mature blood cell comes from a unique hematopoietic stem cell (HSC). Each HSC, however, has tremendous proliferative potential and contributes a large number and variety of mature blood cells for a significant fraction of an animal’s life. Traditionally, HSCs have been viewed as a homogeneous cell population, with each cell possessing equal and unlimited proliferative potential. In other words, the fate of each HSC (to differentiate or replicate) would be determined by its intrinsic stochastic activation and signals from its microenvironment [1, 2].

However, as first shown in Muller-Sieburg *et al.* [3], singly transplanted murine HSCs differ significantly in their long-term lineage (cell-type) output and in their proliferation and differentiation rates [4-7]. Similar findings have been found from examining human embryonic stem cells and HSCs *in vitro* [8, 9]. While cell-level knowledge of HSCs is essential, it does not immediately provide insight into the question of blood homeostasis at the animal level. More concretely, analysis of single-cell transplants does not apply to human bone marrow transplants, which involve millions of CD34-expressing primitive hematopoietic and committed progenitor cells. Polyclonal blood regeneration from such hematopoietic stem and progenitor cell (HSPC) pools is more complex and requires regulation at both the individual cell and system levels to achieve stable [10, 11] or dynamic [12] homeostasis.

To dissect how a population of HSPCs supplies blood, several high-throughput assay systems that can quantitatively track repopulation from an individual stem cell have been developed [6, 11, 13, 14]. In the experiment analyzed in this study, as outlined in Fig. 1, each individual CD34+HSPC is distinctly labeled by the random incorporation of a lentiviral vector in the host genome before transplantation into an animal. All cells that result from proliferation and differentiation of a distinctly marked HSPC will carry identical markings defined by the location of the original viral integration site. By sampling nucleated blood cells and enumerating their unique viral vector integration sites (VIS), one can quantify the cells that arise from a single HSPC marked with a viral vector. Such studies in humans [15] have revealed highly complex polyclonal repopulation that is supported by tens of thousands of different clones [15-18]; a clone is defined as a population of cells of the same lineage, identified here by a unique VIS. These lineages, or clones, can be distributed across all cell types that may be progeny of the originally transplanted HSC after it undergoes proliferation and differentiation. However, the number of cells of any VIS lineage across certain cell types may be different. By comparing abundances of lineages across blood cells of different types, for example, one may be able to determine the heterogeneity or bias of the HSC population or if HSCs often switch their output. This type of analysis remains particularly difficult in human studies since transplants are performed under diseased settings and followed for only 1-2 years.

**Figure 1:**
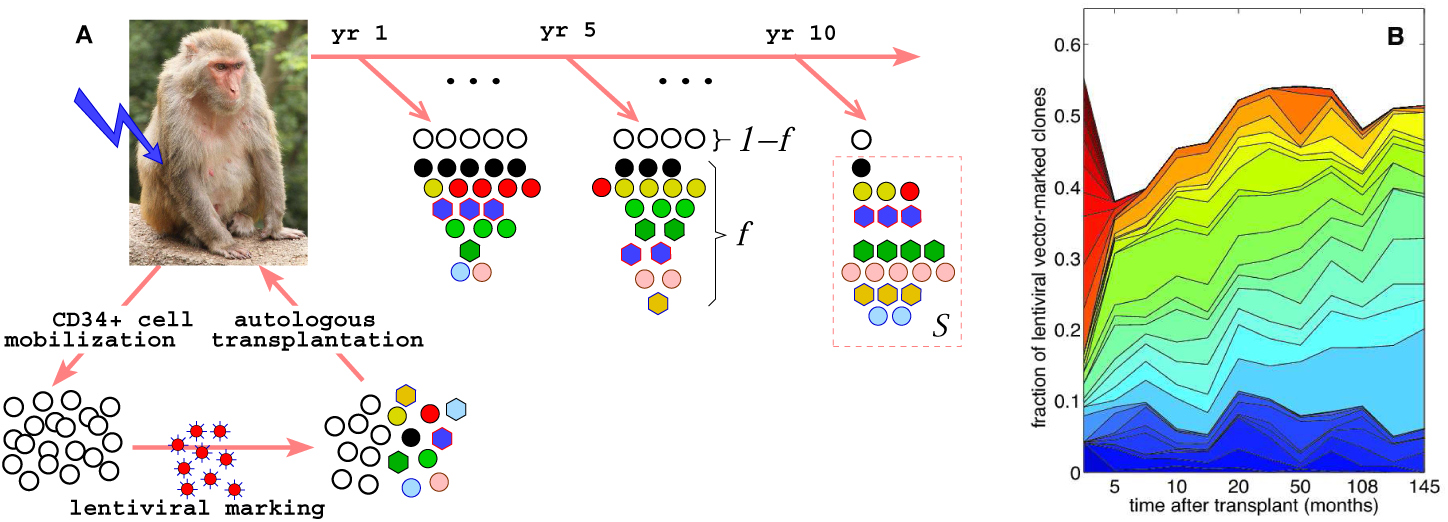
Probing hematopoietic stem and progenitor cell biology (HSPC) through polyclonal analysis. (A) Mobilised CD34+ bone marrow cells from rhesus macaques are first marked individually with lentiviral vectors and transplanted back into the animal after nonlethal myeloablative irradiation [19]. Depending on the animal, 30–160 million CD34+ cells were transplanted, with a fraction ~ 0.07 − 0.3 of them being lentivirus-marked. The clonal contribution of vector-marked HSPCs is measured from blood samples periodically drawn over a dozen years [19]. An average fraction *f* ~ 0.03 − 0.1 of the sampled granulocytes and lymphocytes in the peripheral blood was found to be marked. This fraction is smaller than the fraction of marked CD34+ cells due probably to repopulation by surviving stem cells in the marrow after myeloablative conditioning. Within any post-transplant sample, *S* = 1342 − 44, 415 (average: 10, 026) viral integration sites (VIS) of the marked cells were sequenced (see [14, 19] for details). (B) The fraction of all sequenced VIS reads belonging to each clone is shown by the thickness of the slivers. Small clones are not explicitly shown.

We analyze here a systematic clone-tracking study that used a large number of HSPC clones in a transplant and competitive repopulation setting comparable to that used in humans [19]. In these nonhuman primate (NHP) rhesus macaque experiments, lentiviral vector-marked clones were followed for up to a decade posttransplantation (equivalent to about 30 years in humans if extrapolated by average lifespan). All data are available in the Supplementary Information files of Kim et al. [19]. This long-term study allows one to clearly distinguish HSC clones from other short-term progenitor clones that were included in the initial pool of transplanted CD34+ cells. Hundreds to thousands of detected clones participated in repopulating the blood in a complex yet highly structured fashion. Preliminary examination of some of the clone populations suggests waves of repopulation with short-lived clones that first grow then vanish within the first 1-2 years, depending on the animal [19]. Subsequent waves of HSC clones appear to rise and fall sequentially over the next 4-12 years. This picture is consistent with recent observations in a transplant-free transposon-based tagging study in mice [20] and in human gene therapy [15, 16]. Therefore, the dynamics of clonally tracked NHP HSPC repopulation provide rich data that can inform our understanding of regulation, stability, HSPC heterogeneity, and possibly HSPC aging in hematopoiesis.

Although the time-dependent data from clonal repopulation studies are rich in structure, in this study we focus on one specific aspect of the data: the number of clones that are of a certain abundance as described in Fig. 2. Rather than modeling the highly dynamic populations of each clone, our aim here is to first develop a more global understanding of how the total number of clones represented by specific numbers of cells arises within a mechanistically reasonable model of hematopoiesis. The size distributions of clones present in the blood sampled from different animals at different times are characterised by specific shapes, with the largest clones being a factor of 100–1000 times more abundant than the most rarely detected clones. Significantly, our analysis of renormalised data indicate that the clone size-distribution reaches a stationary state as soon as a few months after transplantation (see Fig. 4 below). To reconcile the observed stationarity of the clone size-distributions with the large diversity of clonal contributions in the context of HSPC-mediated blood repopulation, we developed a mathematical model that treats three distinct cell populations: HSCs, “transit-amplifying” (TA) progenitor cells, and fully differentiated nucleated blood cells (Fig. 3). While multistage models for a detailed description of differentiation have been developed [21], we lump different stages of cell types within the transit-amplifying progenitor pool into one population, avoiding excess numbers of unmeasurable parameters. Another important feature of our model is the overall effect of feedback and regulation, which we incorporate via a population-dependent cell proliferation rate for progenitor cells.

**Figure 2:**
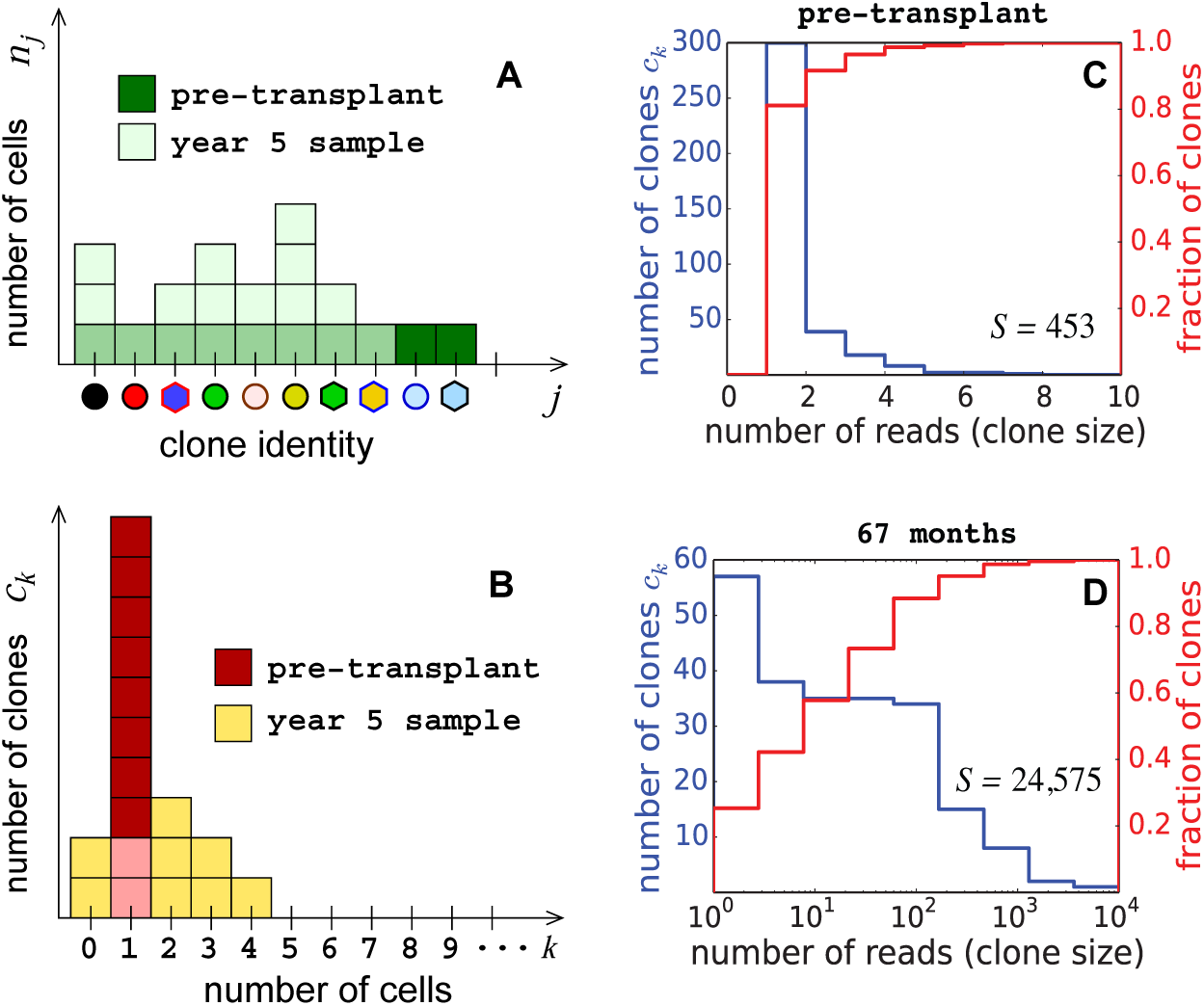
Quantification of marked clones. (A) Assuming each transplanted stem cell is uniquely marked, the initial number of CD34+ cells representing each clone is one. (B) The pre-transplant clone size-distribution is thus defined by the total number of transplanted CD34+ cells and is peaked at one cell. Post-transplant proliferation and differentiation of the HSC clones result in a significantly broader clone size-distribution in the peripheral blood. The number of differentiated cells in each clone and the number of clones represented by exactly *k* cells, five years post-transplantation (corresponding to Fig. 1A), are overlaid in (A) and (B) respectively. (C) Clone size-distribution (blue) and the cumulative normalised clone size-distribution (red) of the pre-transplant CD34+ population. (D) After transplantation, clone size-distributions in the TA and differentiated peripheral cell pools broaden significantly (with clones ranging over four decades in size) but reach a steady-state. The corresponding cumulative normalised distribution is less steep.

The effective proliferation rate will be modeled using a Hill-type suppression that is defined by the limited space for progenitor cells in the bone marrow. Such a regulation term has been used in models of cyclic neutropinia [22] but has not been explicitly treated in models of clone propagation in hematopoiesis. Our mathematical model is described in greater detail in the next section and in the Additional File.

Our model shows that both the large variability and the characteristic shape of the clone size-distribution can result from a slow HSC-to-progenitor differentiation followed by a burst of progenitor growth, both of which are generic features of hematopoietic systems across different organisms. By assuming a homogeneous HSC population and fitting solutions of our model to available data, we show that randomness from stochastic activation and proliferation and a global “carrying capacity” are sufficient to describe the observed clonal structure. We estimate that only a few thousand HSCs may be actively contributing toward blood regeneration at any time. Our model can be readily generalised to include the role of heterogeneity and aging in the transplanted HSCs and provides a framework to quantitatively study physiological perturbations and genetic modifications of the hematopoietic system.

## Models and Methods

Our model explicitly describes three subpopulations of cells: hematopoietic stem cells (HSCs), transit-amplifying (TA) progenitor cells, and terminally differentiated blood cells (see Fig. 3). We will not distinguish between myeloid or lymphoid lineages but will use our model to analyze clone size-distribution data for granulocytes and peripheral blood mononuclear cells (PBMC) independently. Our goal will be to describe how clonal lineages, started from distinguishable HSCs, propagate through the amplification and terminal differentiation processes.

**Figure 3:**
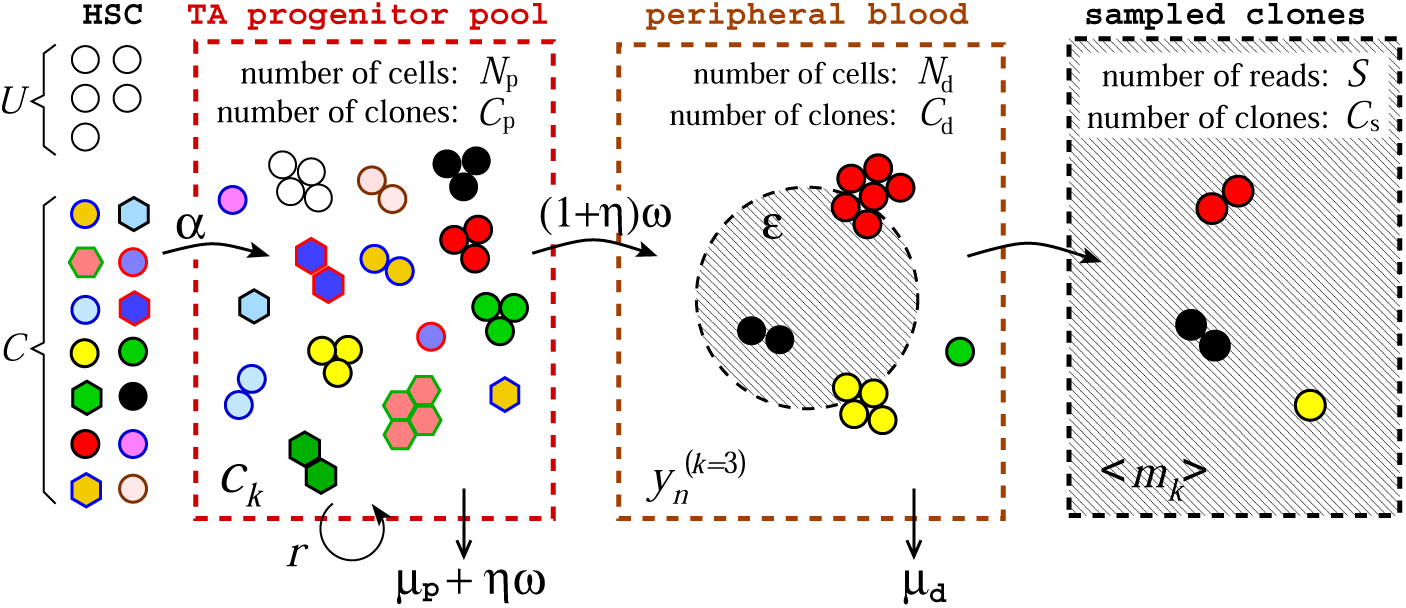
Schematic of our mathematical model. Of the ~ 10^6^ − 10^7^ CD 34+ cells in the animal immediately after transplantation, *C* active HSCs are distinctly labeled through lentiviral vector integration. *U* HSCs are unlabeled because they were not mobilised, escaped lentiviral marking, or survived ablation. All HSCs asymmetrically divide to produce progenitor cells which in turn replicate with an effective, carrying capacity-limited rate *r.* Transit-amplifying progenitor cells die with rate *μ*_p_ or terminally differentiate with rate *ω.* The terminal differentiation of the progenitor cells occurs symmetrically with probability *η* or asymmetrically with probability 1 − *η*. This results in a combined progenitor-cell removal rate *μ* = *μ*_p_ + *ηω*. The differentiated cells outside the bone marrow are assumed not to be subject to direct regulation but undergo turnover with a rate *μ*_d_. The mean total numbers of cells in the progenitor and differentiated populations are denoted *N*_p_ and *N*_d_, respectively. Finally, a small fraction *ε* ≪ 1 of differentiated cells is sampled, sequenced, and found to be marked. In this example, *S* = *εN*_d_ = 5. Because some clones may be lost as cells successively progress from one pool to the next, the total number of clones in each pool must obey *C* ≥ *C*_p_ ≥ *C*_d_ ≥ *C*_s_. Analytic expressions for the expected total number of clones in each subsequent pool are derived in the appendix in the Additional Files.

Often clone populations are modeled directly by dynamical equations for *n_j_* (*t*), the number of cells of a particular clone *j* identified by its specific VIS [23]. Since all cells are identical except for their lentiviral marking, mean-field rate equations for *n_j_*(*t*) are identical for all *j*. Assuming identical initial conditions (one copy of each clone), the expected populations *n_j_*(*t*) would be identical across all clones *j*. This is a consequence of using identical growth and differentiation *rates* to describe the evolution of the *mean* number of cells of each clone.

Therefore, for cells in any specific pool, rather than deriving equations for the mean number *n_j_* of *cells* of each distinct clone *j* (Fig. 2A), we perform a hodograph transformation [24] and formulate the problem in terms of the number of *clones* that are represented by *k* cells, 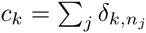 (see Fig. 2B), where the Kronecker *δ*–function 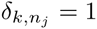 only when *k* = *n_j_* and is zero otherwise. This counting scheme is commonly used in the study of cluster dynamics in nucleation [25] and in other related models describing the dynamics of distributions of cell populations. By tracking the number of clones of different sizes, the intrinsic stochasticity in the *times* of cell division (especially that of the first differentiation event) and the subsequent variability in the clone abundances are quantified. Figures 2A and B qualitatively illustrate *n_j_* and *c_k_*, pre-transplant and after five years, corresponding to the scenario depicted in Fig. 1A. Cells in each of the three pools are depicted in Fig. 3, with different clones grouped according to the number of cells representing each clone.

The first pool (the progenitor cell pool) is fed by HSCs through differentiation. Regulation of HSC differentiation fate is known to be important for efficient repopulation [26, 27] and control [28] and the balance between asymmetric and symmetric differentiation of HSCs has been studied at the microscopic and stochastic levels [29–32]. However, since HSCs have life spans comparable to that of an animal, we reasoned that the total number of HSCs changes only very slowly after the initial few-month transient after transplant. For simplicity, we will assume, consistent with estimates from measurements [33], that HSCs divide only asymmetrically. Therefore, upon differentiation, each HSC produces one partially differentiated progenitor cell and one replacement HSC. How symmetric HSC division might affect the resulting clone sizes is discussed in the Additional File through a specific model of HSC renewal in a finite-sized HSC niche. We find that the incorporation of symmetric division has only a small quantitative effect on the clone size-distribution that we measure and ultimately analyse.

Next, consider the progenitor cell pool. From Fig. 3, we can count the number of clones *c_k_* represented by exactly *k* cells. For example, the black, red, green, and yellow clones are each represented by three cells, so *c*_3_ = 4. Each progenitor cell can further differentiate with rate *ω* into a terminally differentiated cell. If progenitor cells undergo symmetric differentiation with probability *η* and asymmetric differentiation with probability 1 – *η*, the effective rate of differentiation is 2*ηω* + (1 − *η*)*ω* = (1 + *η*)*ω*. In turn, fully differentiated blood cells (not all shown in Fig. 3) are cleared from the peripheral pool at rate *μ*_d_, providing a turnover mechanism. Finally, each measurement is a small-volume sample drawn from the peripheral blood pool, as shown in the final panel in Fig. 3.

Note that the transplanted CD34+ cells contain both true HSCs and progenitor cells. However, we assume that at long times, specific clones derived from progenitor cells die out and that only HSCs contribute to long-lived clones. Since we measure the number of clones of a certain size rather than the dynamics of individual clone numbers, transplanted progenitor cells should not dramatically affect the steady-state clone size-distribution. Therefore, we will ignore transplanted progenitor cells and assume that after transplantation, effectively only *U* unlabeled HSCs and *C* labeled (lentivirus-marked) HSCs are present in the bone marrow and actively asymmetrically differentiating (Fig. 3). Mass-action equations for the expected number of clones *c_k_* of size *k* are derived from considering simple birth-death processes with immigration (HSC differentiation):

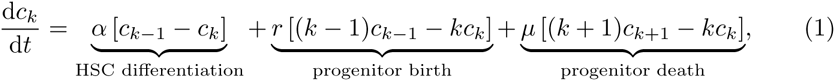

where 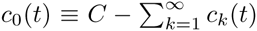 is the number of clones that are not represented in the progenitor pool. We have suppressed the time dependence of *c_k_*(*t*) for notational simplicity. The constant parameter *α* is the asymmetric differentiation rate of all HSCs, while *r* and *μ* are the proliferation and overall clearance rates of progenitor cells. In our model, HSC differentiation events that feed the progenitor pool are implicitly a rate-*α* Poisson process. The appreciable number of detectable clones (Fig. 1B) implies the initial number *C* of HSC clones is large enough that asymmetric differentiation of individual HSCs is uncorrelated. The alternative scenario of a few HSCs undergoing synchronised differentiation would not lead to appreciably different results since the resulting distribution *c_k_* is more sensitive to the progenitor cells’ *unsynchronised* replication and death than to the statistics of the immigration by HSC differentiation.

The final differentiation from progenitor cell to peripheral blood cell can occur through symmetric or asymmetric differentiation, with probabilities *η* and 1 − *η*, respectively. If parent progenitor cells are unaffected after asymmetric terminal differentiation (*i.e.,* dies at its normal rate *μ*_p_), the dynamics are feed-forward and the progenitor population is not influenced by terminal differentiation. Under symmetric differentiation, a net loss of one progenitor cell occurs. Thus, the overall progenitor cell clearance rate can be decomposed as *μ* = *μ*_p_ + *ηω*. We retain the factor *η* in our equations for modeling pedagogy, although in the end it is subsumed in effective parameters and cannot be independently estimated from our data.

The first term in Eqs. 1 corresponds to asymmetric differentiation of each of the *C* active clones, of which *c_k_* are of those lineages with population *k* already represented in the progenitor pool. Differentiation of this subset of clones will add another cell to these specific lineages, reducing *c_k_*. Similarly, differentiation of HSCs in lineages that are represented by *k* − 1 progenitor cells adds cells to these lineages and increase *c_k_*. Note that Eqs. 1 are mean-field rate equations describing the evolution of the expected number of clones of size *k*. Nonetheless, they capture the intrinsic dispersion in lineage sizes that make up the clone size-distribution. While all cells are assumed to be statistically identical, with equal rates *α*, *p*, and *μ*, Eqs. 1 directly model the evolution of the *distribution c_k_* (*t*) that arises ultimately from the distribution of times for each HSC to differentiate or for the progenitor cells to replicate or die. Similar equations have been used to model the evolving distribution of virus capsid sizes [34].

Since the equations for *c_k_*(*t*) describe the evolution of a distribution, they are sometimes described as Master equations for the underlying process [34, 35]. Here we note that the solution to Eqs. 1, *c_k_* (*t*), is the *expected* distribution of clone sizes. Another level of stochasticity could be used to describe the evolution of a *probability distribution* 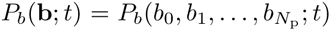 *over the integer numbers b_k_.* This density represents the *probability* that at time *t*, there are *b*_0_ unrepresented lineages, *b*_1_ lineages represented by one cell in the progenitor pool, *b*_2_ lineages represented by two cells in the progenitor pool, and so on. Such a probability distribution would obey an *N*_p_-dimensional Master equation rather than a one-dimensional equation as in Eqs. 1 and once known, can be used to compute the mean 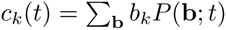. To consider the entire problem stochastically, the variability described by probability distribution *P_b_* would have to be propagated forward to the differentiated cell pool as well. Given the modest number of measured data sets and the large numbers of lineages that are detectable in each, we did not attempt to use the data as samples of the distribution *P_b_* and directly model the mean values *c_k_* instead. Variability from both intrinsic stochasticity and sampling will be discussed in the Additional File.

After defining *u*(*t*) as the number of unlabeled cells in the progenitor pool, and 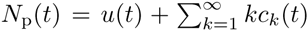 as the total number of progenitor cells, we find 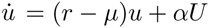 and

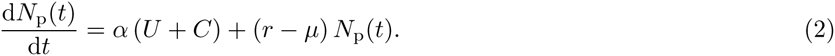

Without regulation, the total population *N*_p_(*t* → ∞) will either reach *N*_p_ ≈ *α*(*U* + *C*)/(*μ*, − *r*) for *μ* > *r* or will exponentially grow without bound for *r* > *μ*. Complex regulation terms have been employed in deterministic models of differentiation [28] and in stochastic models of myeloid/lymphoid population balance [36]. For the purpose of estimating macroscopic clone sizes, we assume regulation of cell replication and/or spatial constraints in the bone marrow can be modeled by a simple effective Hill-type growth law [22, 37]:

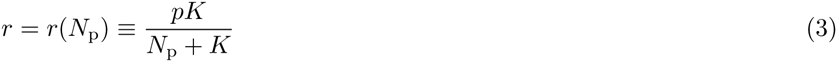

where *p* is the intrinsic replication rate of an isolated progenitor cell. We assume that progenitor cells at low density have an overall positive growth rate *p* > *μ*. The parameter *K* is the cell progenitor cell population in the bone marrow that corresponds to half-maximum of the the effectively growth rate. It can also be interpreted as a limit to the bone marrow size that regulates progenitor cell proliferation to a value determined by *K*, *p*, and *μ* and is analogous to the carrying capacity in logistic models of growth [38]. For simplicity, we will denote *K* as the “carrying capacity” in Eq. 3 as well. Although our data analysis is insensitive to the precise form of regulation used, we chose the Hill-type growth suppression because it avoids negative growth rates that confuse physiological interpretation. An order-of-magnitude estimate of the bone marrow size (or carrying capacity) in the rhesus macaque is *K* ~ 10^9^. Ultimately we are interested in how a limited progenitor pool influences the overall clone size-distribution, and a simple, single-parameter (*K*) approximation to the progenitor-cell growth constraint is sufficient.

Upon substituting the growth law *r*(*N*_p_) described by Eq. 3 into Eq. 2, the total progenitor cell population *N*_p_(*t* → ∞) at long times is explicitly shown in Eq. A19 (see Additional File) to approach a finite value that depends strongly on *K*. Progenitor cells then differentiate to supply peripheral blood at rate (1 + *η*)*ω* so that the total number of differentiated blood cells obeys

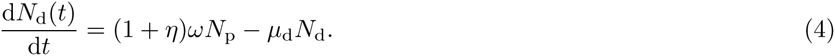

At steady-state, the combined peripheral nucleated blood population is estimated to be *N*_d_ ~ 10^9^ − 10^10^ [39], setting an estimate of *N*_d_/*N*_p_ ≈ (1 + *η*)*ω*/*μ*_d_ ~ 1 − 10. Moreover, as we shall see, the relevant factor in our steady-state analysis will be the numerical *value* of the effective growth rate *r*, rather than its functional form. Therefore, the chosen form for regulation will not play a role in the mathematical results in this paper except to explicitly define parameters (such as *K*) in the regulation function itself.

To distinguish and quantify the clonal structure within the peripheral blood pool, we define 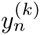 to be the number of clones that are represented by exactly *n* cells in the differentiated pool *and k* cells in the progenitor pool. For example, in the peripheral blood pool shown in Fig. 3, 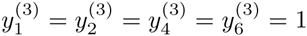. This counting of clones across both the progenitor and peripheral blood pools is necessary in order to balance progenitor cell differentiation rates with peripheral blood turnover rates. The evolution equations for 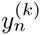 can be expressed as

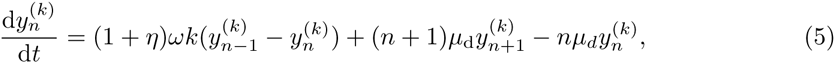

where 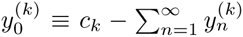 represents the number of progenitor clones of size *k* that have not yet contributed to peripheral blood. The transfer of clones from the progenitor population to the differentiated pool arises through 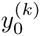 and is simply a statement that the number of clones in the peripheral blood can increase only by differentiation of a progenitor cell whose lineage has not yet populated the peripheral pool. The first two terms on the right-hand side of Eq. 5 represent immigration of clones represented by *n* − 1 and *n* differentiated cells *conditioned upon* immigration from only those specific clones represented by *k* cells in the progenitor pool. The overall rate of addition of clones from the progenitor pool is thus (1 + *η*)*ωk*, in which the frequency of terminal differentiation is weighted by the stochastic division factor (1 + *η*). By using the Hill-type growth term *r*(*N*_p_) from Eq. 3, Eq. 1 can be solved to find *c_k_* (*t*), which in turn can be used in Eq. 5 to find 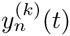. The number of clones in the peripheral blood represented by exactly *n* differentiated cells is thus 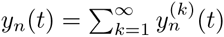.

As we mentioned, Eqs. 1 and 5 describe the evolution of the expected clone size-distribution. Since each measurement represents one realization of the distributions *c_k_*(*t*) and *y_n_*(*t*), the validity of Eqs. 1 and 5 relies on a sufficiently large *C* such that the marked HSCs generate enough lineages and cells as to adequately sample the subsequent peripheral blood clone size-distribution. In other words, measurement-to-measurement variability described by *e.g.,* 〈*c_k_*(*t*)*c_k_*′(*t*)〉 − 〈*c_k_*(*t*)〉〈*c_k_*′(*t*)〉 is assumed negligible (see Additional File). Our modeling approach would not be applicable to studying single HSC transplant studies [4–6] unless the measured clone sizes from multiple experiments are aggregated into a distribution.

Finally, in order to compare model results with animal blood data, we must consider the final step of sampling small aliquots of the differentiated blood. As derived in Eq. A11 of the Additional File, if *S* marked cells are drawn and sequenced successfully (from a total differentiated cell population *N*_d_), the expected number of clones 〈*m_k_*(*t*)〉 represented by *k* cells is given by

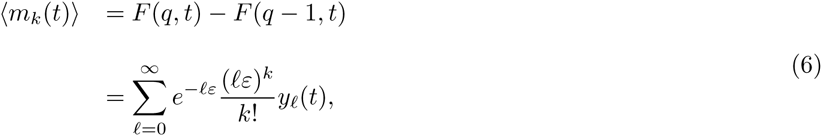

where *ε* ≡ *S/N*_d_ ≪ 1 and 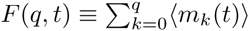 is the sampled, expected cumulative size-distribution. Upon further normalization by the total number of detected clones in the sample, *C_s_*(*t*) = *F*(*S*, *t*) − *F* (0, *t*), we define

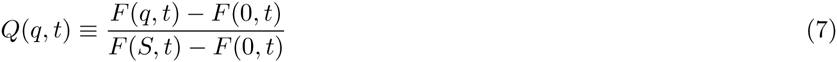

as the fraction of the total number of sampled clones that are represented by *q* or fewer cells. Since the data represented in terms of *Q* will be seen to be time-independent, explicit expressions for 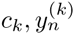, 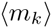, and *Q*(*q*) can be derived. Summarizing, the main features and assumptions used in our modeling include:

- a neutral-model framework [40] that directly describes the distribution of clone sizes in each of the four cell pools: HSCs, progenitor cells, peripheral blood cells, and sampled blood cells. The cells in each pool are statistically identical.
- a constant, asymmetric HSC differentiation rate *α*. The appreciable numbers of unsynchronised HSCs allow the assumption of Poisson-distributed differentiation times of the HSC population. The level of differentiation symmetry is found to have little effect on the steady-state clone size-distribution (see Additional File). Symmetry of the terminal differentiation step is also irrelevant for understanding the available data.
- a simple one-parameter (*K*) growth regulation model that qualitatively describes the finite maximum size of the progenitor population in the bone marrow. Ultimately, the specific form for the regulation is unimportant since only the steady-state value of the growth parameter *r* affects the parameter fitting.

Using only these reasonable model features, we are able to compute clone size-distributions and compare them with data. An explicit form for the expected steady-state clone size-distribution 〈*m_k_*〉 is given in Eq. A32 of the Additional File, and the parameters and variables used in our analysis are listed in Table 1.

**Table 1:** Table of model parameters and variables. Estimates of steady-state values are provided where available. We assume little prior knowledge on all but a few of the more established parameters. Nonetheless, our modeling and analysis places constraints on combinations of parameters, allowing us to fit data and provide estimates for steady-state values of *U* + *C* ~ 10^3^ − 10^4^ and *α*(*N*_p_ + *K*)/(*pK*) ~ 0.002 − 0.1.

| symbol | parameter or variable | estimate | Ref. |
| --- | --- | --- | --- |
| $\alpha$ | single HSC asymmetric differentiation rate | $\sim 0.1 - 0.3/\text{mo}$ | [46, 51] |
| $p$ | free progenitor cell proliferation rate | | |
| $\mu_p$ | progenitor cell death rate | | |
| $\mu_d$ | differentiated cell death rate | $\sim 0.01 - 0.3/\text{day}$ | [52, 53] |
| $\eta$ | symmetric differentiation probability | | |
| $\omega$ | terminal differentiation rate | | |
| $K$ | progenitor cell capacity | $\sim 10^9$ | |
| $U$ | number of active unmarked HSCs | | |
| $C$ | number of active viral-marked HSCs | | |
| $C_p$ | number of progenitor clones | | Eq. A23 |
| $C_d$ | number of differentiated clones | | Eq. A23 |
| $C_s$ | number of sampled clones | | Eq. A24 |
| $S$ | number of sequences read | $\sim 10^3 - 10^4$ | [14] |
| $c_k$ | number of progenitor clones of size $k$ | | |
| $u$ | unlabelled progenitor cell population | | |
| $N_p$ | total progenitor cell population | | |
| $N_d$ | total differentiated blood population | $\sim 10^9 - 10^{10}$ | |
| $y_n^{(k)}$ | number of differentiated clones of size $n$ arising from progenitors of size $k$ | | |

## Results and Analysis

In this section, we describe how previously published data (available in the Supplementary Information files of Kim et al. [19])—the number of cells of each detected clone in a sample of the peripheral blood—are used to constrain parameter values in our model. We emphasize that our model is structurally different from models used to track lineages and clone size-distributions in retinal and epithelial tissues [41, 42]. Rather than tracking only the lineages of stem cells—which are allowed to undergo asymmetric differentiation, symmetric differentiation, or symmetric replication—our model assumes a highly proliferative population constrained by a carrying capacity *K* and slowly fed at rate α by an asymmetrically dividing HSC cell pool of *C* fixed clones. We have also included terminal differentiation into peripheral blood and the effects of sampling on the expected clone size-distribution. These ingredients yield a clone size-distribution different from those previously derived [41, 42], as described in more detail below.

**Stationarity in time.** Clonal contributions of the initially transplanted HSC population have been measured over 4–12 years in four different animals. As depicted in Fig. 4A, populations of individual clones of peripheral blood mononuclear cells (PBMC) from animal RQ5427, as well as all other animals, show significant variation in their dynamics. Since cells of any detectable lineage will number in the millions, this variability in lineage size across time cannot be accounted for by the intrinsic stochasticity of progenitor cell birth and death. Rather, these rises and falls of lineages likely arises from a complicated regulation of HSC differentiation and lineage aging. However, in our model and analysis, we do not keep track of lineage sizes *n_i_*. Instead, define *Q*(*v*) as the fraction of clones arising with relative frequency *v* = *fq*/*S* or less (here, *q* is the number of VIS reads of any particular clone in the sample, *f* is the fraction of all sampled cells that are marked, and *S* is the total number of sequencing reads of marked cells in a sample). Fig. 4B shows data analyzed in this way and reveals that *Q*(*v*) appears stationary in time.

**Figure 4:**
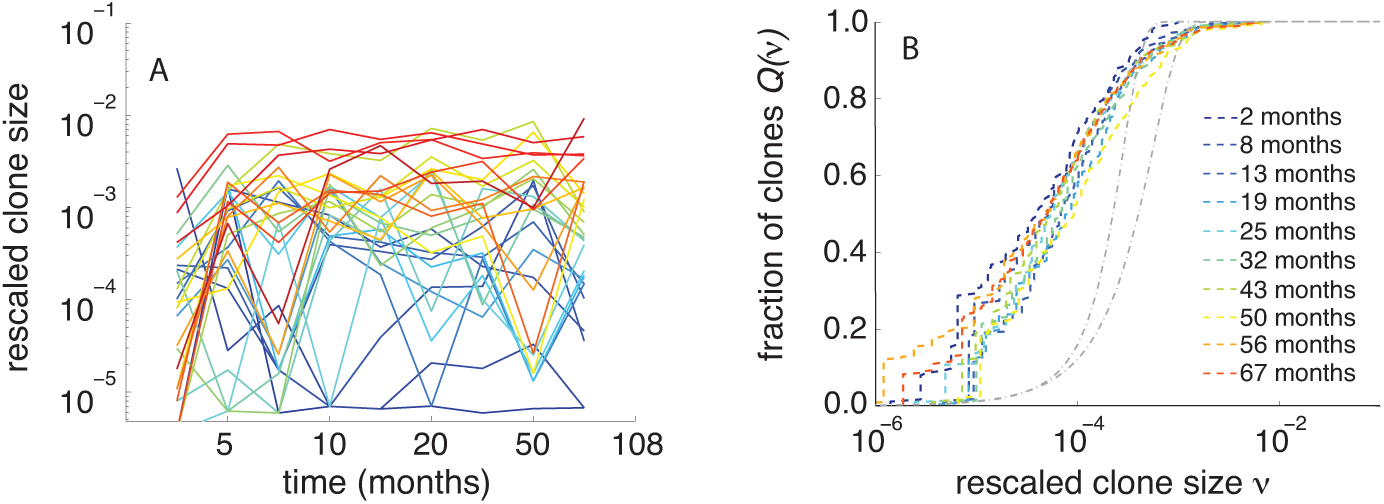
(A) Individual clone populations (here, PBMC of animal RQ5427) show significant fluctuations in time. For clarity, only clones that reach an appreciable frequency are plotted. (B) The corresponding normalised clone size-distributions at each time point are rescaled by the sampled and marked fraction of blood, 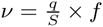, where *q* is the number of reads of a particular clone within the sample. After an initial transient, the fraction of clones (dashed curves) as a function of relative size remains stable over many years. For comparison, the dot-dashed grey curves represent binomial distributions (with *S* = 10^3^ and 10^4^ and equivalent mean clone sizes) and underestimate low population clones.

The observed steady-state clone size-distribution is broad, consistent with the mathematical model developed above. The handful of most populated clones constitutes up to 1–5% of the entire differentiated blood population. These dominant clones are followed by a large number of clones with fewer cells. The smallest clones sampled in our experiment correspond to a single read *q* = 1, which yields a minimum measured frequency *v*_min_ = *f/S*. A single read may comprise only 10^−4^ − 10^−3^% of all differentiated blood cells. Note that the cumulative distribution *Q*(*v*) exhibits higher variability at small sizes simply because fewer clones lie below these smaller sizes.

Although engraftment occurs within a few weeks and total blood populations *N*_p_ and *N*_d_ (and often immune function) re-establish themselves within a few months after successful HSC transplant [43, 44], it is still surprising that the clone size-distribution is relatively static within each animal (see Additional File for other animals). Given the observed stationarity, we will use the steady-state results of our mathematical model (explicitly derived in the Additional File) for fitting data from each animal.

### Implications and model predictions

By using the exact steady-state solution for *c_k_* (Eq. A21) in Eq. A18 of the Additional File, we can explicitly evaluate the expected clone size-distribution 〈*m_k_*〉 using Eq. 6, and the expected cumulative clone fraction *Q*(*q*) using Eq. 7. In steady-state, the clone size-distribution of progenitor cells can also be approximated by a Gamma distribution with parameters *a* ≡ *α*/*r* and 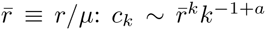 (see Eq. A27 in the Additional File). In realistic steady-state scenarios near carrying capacity, 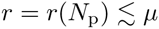, as calculated explicitly in Eq. A20 of the Additional File. By defining 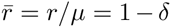, we find that *δ* is inversely proportional to the carrying capacity:

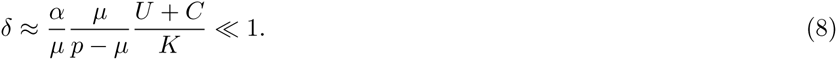

The dependences of 〈*m_q_*〉 on *δ* and *a* = *α*/*r* are displayed in Fig. 5A, in which we have defined *w* ≡ (1 + *η*)*ω*/*μ*_d_.

**Figure 5:**
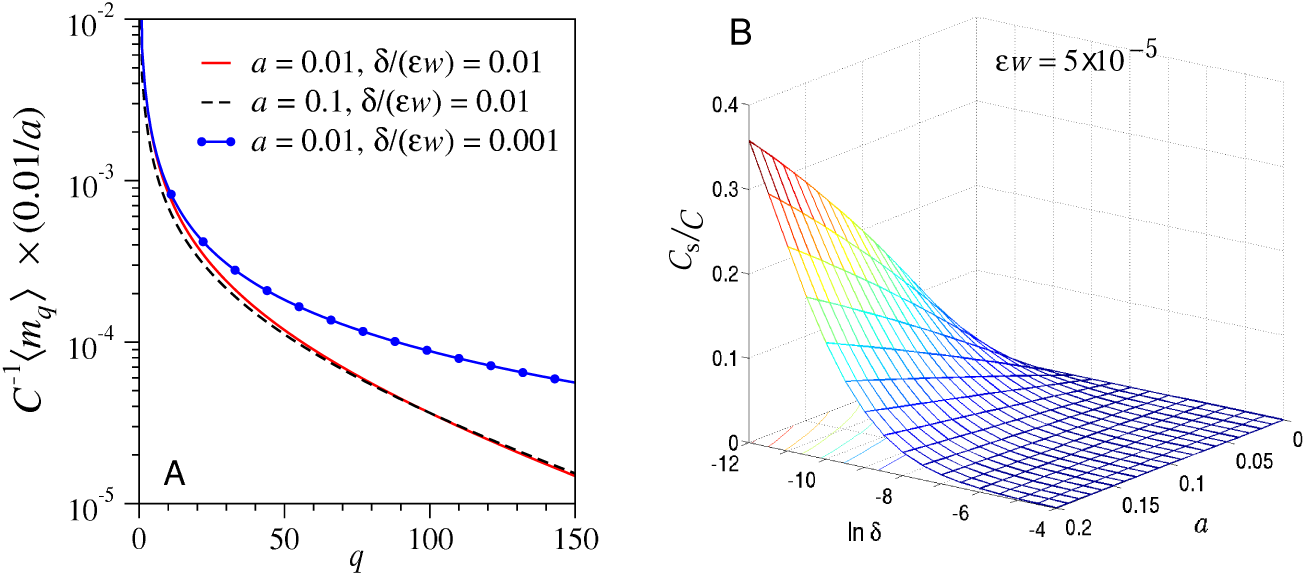
(A) Expected clone size-distributions *C*^−1^〈*m_q_*〉 derived from the approximation in Eq. A32 are plotted for various *a* and *δ/*(*εw*) (where *w* ≡ (1 + *η*)*ω/μ*_d_). The nearly coincident solid and dashed curves indicate that variations in *a* mainly scale the distribution by a multiplicative factor. In contrast, the combination *δ/*(*εw*) controls the weighting at large clone sizes through the population cut-off imposed by the carrying capacity. Of the two controlling parameters, the steady-state clone size-distribution is most sensitive to *R* ≅ *δ/*(*εw*). The dependence of data-fitting on these two parameters is derived in the Additional File and discussed in the next section. (B) For *εw* = 5 × 10^−5^, the expected fraction *C*_s_/*C* of active clones sampled as a function of ln *δ* and *α.* The expected number of clones sampled increases with carrying capacity *K*, HSC differentiation rate *a = α/r*, and the combined sampling and terminal differentiation rate *εw.*

Although our equations form a mean-field model for the expected number of measured clones of any given size, randomness resulting from the stochastic differentiation times of individual HSCs (all with the same rate α) are taken into account. This is shown in the Additional File (Eqs. A36-A39) where we explicitly consider the fully stochastic population of a single progenitor clone that results from the differentiation of a single HSC. Since independent, unsynchronised HSCs differentiate at times that are exponentially distributed (with rate *α*), we construct the expected clone size-distribution by from this birth-death-immigration process [45] to find a result equivalent to that derived from our original model (Eqs. 1 and A21). Thus, we conclude that if *a* = *α*/*r* is small, the shape of the expected clone size-distribution is mainly determined at short times by the initial repopulation of the progenitor cell pool.

Our model also suggests that the expected number of sampled clones relative to the number of active transplanted clones (see Eq. A24) can be expressed as

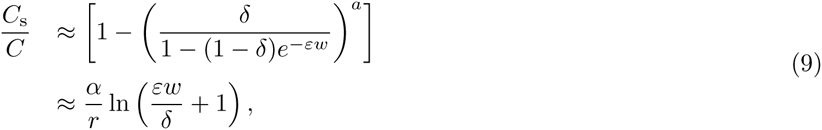

where the last approximation is accurate for *εw* ≪ 1 and *C*_s_/*C* ≪ 1. This clonal diversity one expects to measure in the peripheral blood sample is sensitive to the combination of biologically relevant parameters and rates *δ* and *a* = *α*/*r*. Fig. 5B shows the explicit dependence of the fraction of active clones on *a* and the combination of parameters defining *δ*, for *εw* = *ε*(1 + *η*)*ω*/*u*_d_ = 5 × 10^−5^.

Our analysis shows how scaled measurable quantities such as *C*_s_/*C* and *C*^−1^ 〈*m_q_*〉 depend on just a few combinations of experimental and biological parameters. This small domain of parameter sensitivity reduces the number of parameters that can be independently extracted from clone size-distribution data. For example, the mode of terminal differentiation described by *η* clearly cannot be extracted from clonal tracking measurements. Similarly, more detailed models of the complex regulation processes would introduce additional parameters that are not resolved by these experiments. Nonetheless, we shall fit our data and known information contained in the experimental protocol to our model to estimate biologically relevant parameters such as the total number of activated HSCs *U* + *C*, and thus indirectly *C*.

### Model fitting

Our mathematical model for 〈*m_k_*〉 (and *F*(*q*) and *Q*(*q*)) includes numerous parameters associated with the processes of HSC differentiation, progenitor cell amplification, progenitor cell differentiation, peripheral blood turnover, and sampling. Data fitting is performed using clone size-distributions derived separately from the read counts from both the left and right ends of each VIS (see [14] for details on sequencing). Even though we fit our data to 〈*m_k_*〉 using three independent parameters *a* = *α*/*r*, 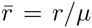, and *εw*, we found that within the relevant physiological regime, all clone distributions calculated from our model are most sensitive to just two combinations of parameters (see Additional File for an explicit derivation):

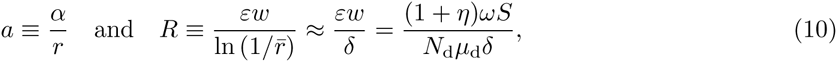

where the last approximation for *R* is valid when 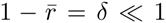. While the fits are rather insensitive to *εw* this parameter can fortunately be approximated from estimates of *S* and the typical turnover rate of differentiated blood. Consequently, we find two maximum likelihood estimates (MLEs) for *a* and *R* at each time point. It is important to note that fitting our model to steady-state clone size-distributions does not determine all of the physiological parameters arising in our equations. Rather, they provide only two constraints that allow one to relate their values.

For ease of presentation, we henceforth show all data and comparison with our model equations in terms of the fraction *Q*(*v*) or *Q*(*q*) (Fig. 4B or 6A, B). Figs. 6A, B shows MLE fitting to the raw data 〈*m_k_*〉 plotted in terms of the normalised but unrescaled data *Q*(*q*) for two different peripheral blood cell types from two animals (RQ5427 and RQ3570). Data from all other animal are shown and fitted in the Additional File, along with overall goodness-of-fit metrics. Raw cell count data were given in Kim *et al.* [19].

**Figure 6:**
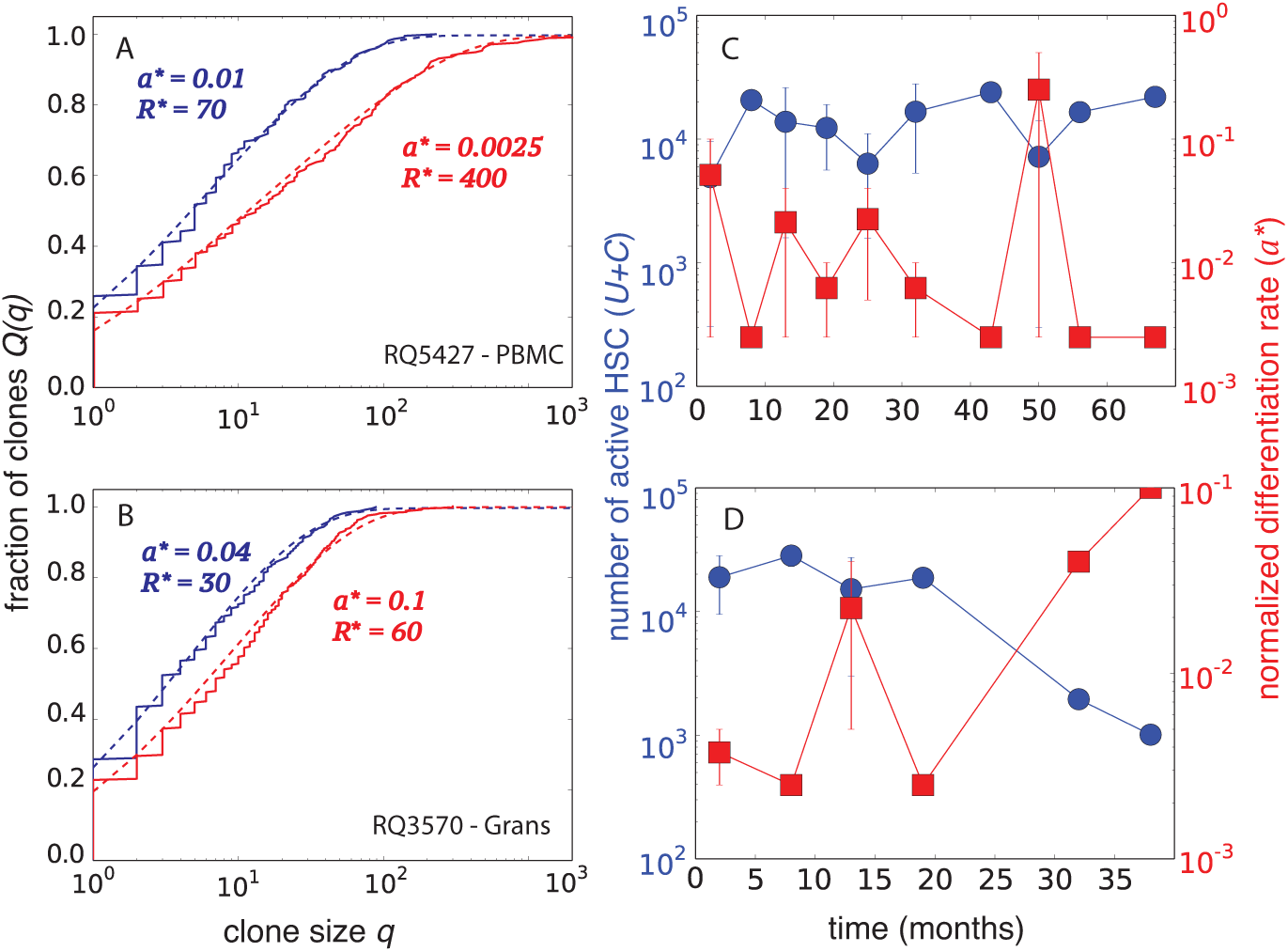
(A) Fitting raw (not rescaled, as shown in Fig. 4) clone size-distribution data, to 〈*m_k_*〉 from Eq. 6 at two time points for animal RQ5427. The maximum likelihood estimates (MLEs) are (*a*\* ≈ 0.01, *R*^*^ ≈ 70) and (*a*^*^ ≈ 0.0025, *R*^*^ ≈ 400) for data, taken at 32 (blue) and 67 (red) months post-transplant, respectively. Note that the MLE values for different samples vary primarily due to different values of *S* (and hence *ε*) used in each measurement. (B) For animal RQ3570, the clone fractions at 32 (blue) and 38 (red) months yield (*a*^*^ ≈ 0.04, *R*^*^ ≈ 30) and (*a*^*^ ≈ 0.1, *R*^*^ ≈ 60), respectively. For clarity, we show the data and fitted models in terms of *Q*(*q*). (C) Estimated number of HSCs *U+C* (circles) and normalised differentiation rate *a* (squares) for animal RQ5427. (D) *U* + *C* and *a* for animal RQ3570. Note the temporal variability (but also long-term stability) in the estimated number of contributing HSCs. Additional details and fits for other animals are qualitatively similar and given in the Additional File.

### HSC asymmetric differentiation rate

The MLE for *a* = *α*/*r*, *a*^*^, was typically in the range 10^−2^ − 10^−1^. Given realistic parameter values, this quantity mostly provides an estimate of the HSC relative differentiation rate *a*^*^ ~ *α*/(*μ*_p_ + *ηω*). The smallness of *a*^*^ indicates a slow HSC differentiation relative to the progenitor turnover rate *μ*_p_ and the final differentiation rate *ηω*, consistent with the dominant role of progenitor cells in populating the total blood tissue. Note that besides the intrinsic insensitivity to *εw*, the goodness-of-fit is also somewhat insensitive to small values of *a*^*^ due to the weak dependence of *c_k_* ~ 1/*k*^1−^*^a^* on *a* (see Additional File). The normalised relative differentiation rates estimated from two animals are shown by the squares (right axis) in Fig. 6C,D.

**Number of HSCs.** The stability of blood repopulation kinetics is also reflected in the number of estimated HSCs that contribute to blood (shown in Fig. 6C,D). The total number of HSCs is estimated by expressing *U* + *C* in terms of the effective parameters, *R* and *a*, which in turn are functions of microscopic parameters (*α*, *p*, *μ*_p_, *μ*_d_, *w*, *K*) that cannot be directly measured. In the limit of small sample size, *S* ≪ *R*^*^*K*; however, we find *U* + *C* ≈ *S*/(*R*^*^*a*^*^) (see Additional Files), which can then be estimated by using the MLEs *a*^*^ and *R*^*^ obtained by fitting clone size-distributions. The corresponding values of *U* + *C* for two animals are shown by the circles (left axis) in Fig. 6C,D. Although variability in the MLEs exists, the fluctuations appear stationary over the course of the experiment for each animal (see Additional File).

## Summary and Conclusions

Our clonal tracking analysis revealed that individual clones of HSCs contributed differently to the final differentiated blood pool in rhesus macaques, consistent with mouse and human data. Carefully replotting the raw data (clone sizes) in terms of the normalised, rescaled cumulative clone size-distribution (the fraction of all detected clones that are of a certain size or less) shows that these distributions reach steady-state a few months after transplantation. Our results carry important implications for stem cell biology. Maintaining homeostasis of the blood is a critical function for an organism. Following myeloablative stem cell transplant, the hematopoietic system must repopulate rapidly to ensure survival of the host. Not only do individual clones rise and fall temporally, as previously shown [19], but as any individual clone of a certain frequency declines, it is replaced by another of similar frequency. This “exchange-correlated” mechanism of clone replacement may provide a mechanism by which overall homeostasis of hematopoiesis is maintained long-term, thus ensuring continued health of the blood system.

To understand these observed features and the underlying mechanisms of stem cell-mediated blood regeneration, we developed a simple neutral population model of the hematopoietic system that quantifies the dynamics of three subpopulations: hematopoietic stem cells (HSCs), transit-amplifying (TA) progenitor cells, and fully differentiated nucleated blood cells. We also include the effects of global regulation by assuming a Hill-type growth rate for progenitor cells in the bone marrow but ignore cell-to-cell variation in differentiation and proliferation rates of all cells.

Even though we do not include possible HSC heterogeneity, variation in HSC activation, progenitor cell regulation, HSC and progenitor-cell ageing (“progenitor bursting”), niche- or signal molecule-mediated controls, or intrinsic genetic/epigenetic differences, solutions to our simple *homogeneous* HSC model are remarkably consistent with observed clone size-distributions. As a first step, we focus on how the intrinsic stochasticity in just the cellular birth, death, and differentiation events gives rise to the progenitor clone size-distribution.

To a large extent, the exponentially distributed first HSC differentiation times and the growth and turnover of the progenitor pool control the shape of the expected long-time clone size-distribution. Upon constraining our model to the physiological regime relevant to the experiments, we find that the calculated shapes of the clone size-distributions are sensitive to effectively only two composite parameters. The HSC differentiation rate *α* sets the scale of the expected clone size-distribution but has little effect on the shape. Parameters including carrying capacity *K*, active HSCs *U* + *C*, and birth/death rates *p*, *ω*, *μ*_p_, *μ*_d_ influence the shape of the expected clone size-distribution 〈*m_q_*〉 only through the combination *R*, and only at large clone sizes.

Our analysis allowed us to quantitatively estimate other combinations of model parameters. Using a maximum likelihood estimate (MLE), we find values for the effective HSC differentiation rate *a*\* ~ 10^−2^ − 10^−1^ and the number of HSCs that are contributing to blood within any given time frame *U* + *C* ~ 10^3^–10^4^. Since the portion of HSCs that contribute to blood may be varying across their typical lifespan *L* ~ 25yr, the total number of HSCs can be estimated by (*U* + *C*) × *L/τ*, where *τ* ~ 1 yr [19]. Our estimate of a total count of ~ 3 × 10^4^ − 3 × 10^5^ HSCs is about 30-fold higher than the estimate of Abkowitz *et al.* [33] but is consistent with Kim *et al.* [19]. Note that the ratio of *C* to the total number of initially transplanted CD34+ cells provides a measure of the overall potency of the transplant towards blood regeneration. In the extreme case in which one HSC is significantly more potent (through, *e.g.,* a faster differentiation rate), this ratio would be smaller. An example of this type of heterogeneity would be an HSC with one or more cancer-associated mutations, allowing it to out-compete other transplanted normal HSCs. Hence, our clonal studies and the associated mathematical analysis can provide a framework for characterizing normal clonal diversity as well as deviations from it, which may provide a metric for early detection of cancer and other related pathologies.

Several simplifying assumptions have been invoked in our analysis. Crucially, we assumed HSCs divided only asymmetrically and ignored instances of symmetric self-renewal or symmetric differentiation. The effects of symmetric HSC division can be quantified in the steady-state limit. In previous studies, the self-renewal rate for HSCs in primates is estimated between 4–9 months [46, 47], which is slightly longer than the short time scale (~2–4 months) on which we observe stabilization of the clone size-distribution. Therefore, if the HSC population slowly increases in time through occasional symmetric division, the clone size-distribution in the peripheral blood will also shift over long times. The static nature of the clone distributions over many years suggests that size distributions are primarily governed by mechanisms operating at shorter time scales in the progenitor pool. For an HSC population (such as cancerous or precancerous stem cells [48]) that has already expanded through early replication, the initial clone size-distribution within the HSC pool can be quantified by assuming an HSC pool with separate carrying capacity KHSC. Such an assumption is consistent with other analyses of HSC renewal [49]. All our results can be used (with the replacement *C* → *K*_HSC_) if the number of transplanted clones *C* ≥ *K*_HSC_ because replication is suppressed in this limit. When *K*_HSC_ ≫ *C* ≫ 1, replicative expansion generates a broader initial HSC clone size-distribution (see Additional File). The resulting final peripheral blood clone size-distribution can still be approximated by our result (Eq. 6) if the normalised differentiation rate *a* ≪ 1, exhibiting the insensitivity of the differentiated clone size-distribution to a broadened clone size-distribution at the HSC level. However, if HSC differentiation is sufficiently fast 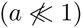, the clonal distribution in the progenitor and differentiated pools can be modified.

To understand temporal dynamics of clone size-distributions, a more detailed numerical study of our full time-dependent neutral model is required. Such an analysis can be used to investigate the effects rapid temporal changes in HSC division mode [41]. Temporal models would also allow investigation into the evolution of HSC mutations and help unify concepts of clonal stability (as indicated by the stationarity of rescaled clone size-distributions) with ideas of clonal succession [10, 11] or dynamic repetition [12] (as indicated by the temporal fluctuations in the estimated number *U* + *C* of active HSCs). Predictions of the time-dependent behavior of clone size-distributions will also prove useful in guiding future experiments in which the animals are physiologically perturbed via *e.g.,* myeloablation, hypoxiation, and/or bleeding. In such other experimental settings, regulation may also occur at the level of HSC differentiation (*α*) and a different mathematical model may be more appropriate.

We have not addressed the temporal fluctuations in *individual* clone abundances evident in our data (Fig. 4A) or in the wave-like behavior suggested by previous studies [19]. Since the numbers of detectable cells of each VIS lineage in the whole animal are large, we believe these fluctuations do not arise from intrinsic cellular stochasticity or sampling. Rather, they likely reflect slow timescale HSC transitions between quiescent and active states and/or HSC ageing [50]. Finally, subpopulations of HSCs that have different intrinsic rates of proliferation, differentiation, or clearance can then be explicitly treated. As long as each subtype in a heterogeneous HSC or progenitor cell population does not convert into another subtype, the overall aggregated clone size-distribution 〈*m_k_*〉 will preserve its shape. Although steady-state data is insufficient to provide resolution of cell heterogeneity, more resolved temporal data may allow one to resolve different parameters associated with different cell types. Such extensions will allow us to study the temporal dynamics of individual clones and clone populations in the context of cancer stem cells and will be the subject of future work.

## Authors’ contributions

TC and SG designed and developed the mathematical modeling and data analysis; TC, SG, and SK wrote the manuscript; SK and IC participated in study design and data interpretation.

## Acknowledgements

This work was supported by grants from the NIH (R01AI110297 and K99HL116234), the California Institute of Regenerative Medicine (TRX-01431), the UCLA AIDS Institute/Center for AIDS Research (AI28697), the NSF PHY11-25915 (KITP/UCSB), and the Army Research Office (W911NF-14-1-0472). The authors also thank B. Shraiman and R. K. P. Zia for helpful discussions.

## Additional Files: Mathematical Appendices and Data Fitting

Here, we provide additional details of our model, including explicit derivations of the sampled clone size distribution, clone size distributions in the steady-state limit, and the effective parameters that accurately describe our data. We also describe the maximum likelihood estimation used to estimate these parameters.

Derivation of sampled clone size distribution:

We first derive an expression for the expected clone size distribution 〈*m_k_*(*t*)〉 in a sample of the differentiated blood, as given by Eq. 6. Define *s_jℓ_* to be the number of cells sampled from the *j*^th^ clone of those that are represented by *ℓ* cells. At any time, the probability that the configuration *s_jℓ_* is observed in a sample of *S* cells can be written

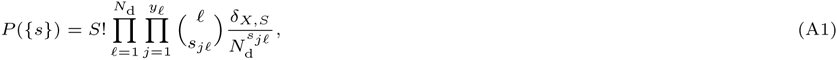

where 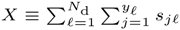, the factor 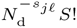 represents the probability that *s_jℓ_* cells were drawn from the *N*_d_ within a sample of *S* cells, and 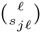 is the number of ways of drawing *s_jℓ_* cells. Finally, the last Kronecker *δ*–function forces the sum over all *s_jℓ_* to equal the total number of cells sampled and sequenced. In any particular sample, the number of clones with size *k* is exactly

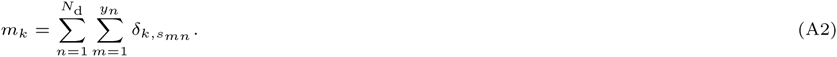

The expected value of this quantity is

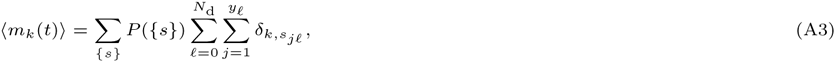

which can be found by using the generating function 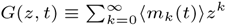:

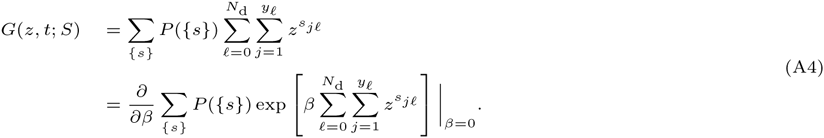

After using the Fourier representation of the Kronecker *δ*—function in Eq. A1, 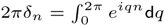, we can further reduce the generating function to

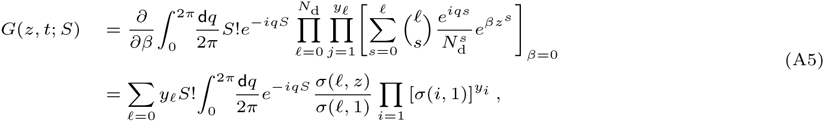

where

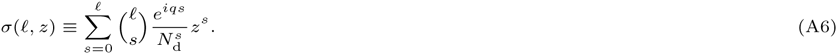

Note that *σ*(*ℓ*, *z*) ≡ (1 + *ze^iq^*/*N*_d_)*^ℓ^*, and that 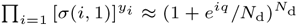. Since *N*_d_ ≫ *S* ≫ 1, and *N*_d_ ~ 10^9^ − 10^10^, we can take the large *N*_d_ limit before the large *S* limit to find 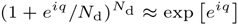, 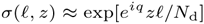, and

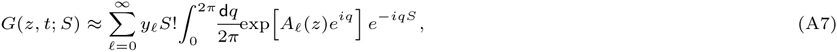

where *A_ℓ_*(*z*) = 1 + *ℓ*(*z* − 1)/*N*_d_. Note that the integral is simply Euler’s integral for 1/Γ(*S* + 1). Namely, we find

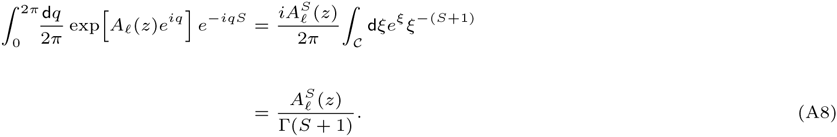

Since 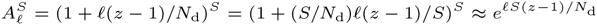 for *S* ≫ 1, we find, for *ε* = *S/N*_d_ ≪ 1,

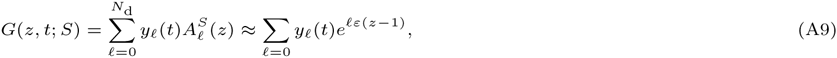

Next, we define the fraction of clones of size 1 ≤ *q* ≤ *S* or less. This distribution includes unrepresented or lost clones, and is defined as 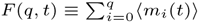. By using Eq. A9 and the definition of *G*(*z*, *t*), we find

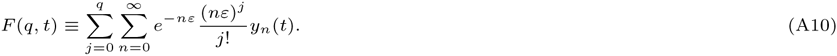

The expected clone size distribution is thus defined as

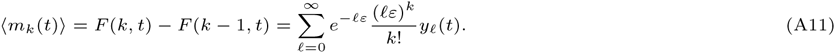

In general, further development of *F*(*k*, *t*) and (*m_k_*(*t*)) requires numerical solution of *c_k_* (*t*) and 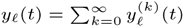. The time-dependence of *F*(*q*, *t*) is further complicated by the time-dependence of *ε*(*t*) = *S*/*N*_d_(*t*), requiring the solution to Eq. 4.

The variability of *m_k_* due to sampling can be also estimated by calculating 〈*m_k_m_k_*_′_〉, which we write as

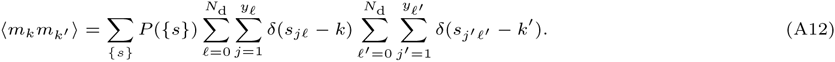

This calculation requires evaluation of the two-dimensional generating function

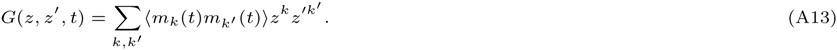

After using Eq. A1 for *P*({*s*}) in Eq. A12 and performing some algebra, we find

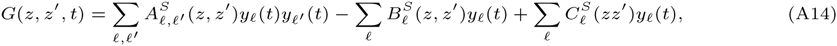

where

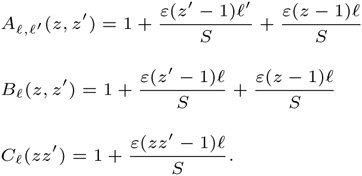

Using (1 + *x*/*S*)*^S^* ≈ *e^x^*, and expanding in powers of *z* and *z*′, we find

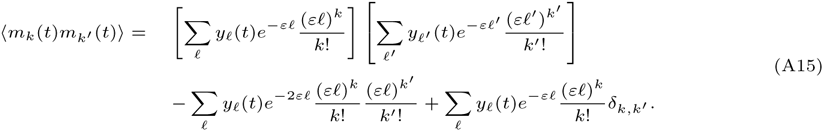

The diagonal variance is simply

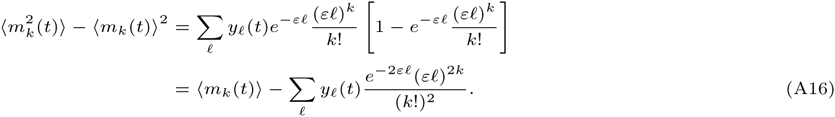

The second term is much smaller than the first except for very small values of *k*. Therefore, the relative fluctuation in *m_k_* due to sampling is

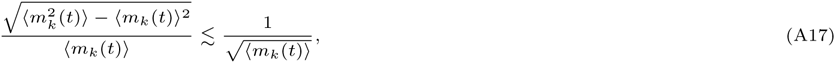

explicitly indicating that the relative fluctuations in the measured number of clones of size *k* decreases as the square-root of its expected value.

Steady-state solution:

As was discussed, the total peripheral blood population in the animals recovered quickly, usually within a few weeks after transplantation. Moreover, from our data, the overall qualitative shape of the clone size distribution also reaches steady-state only after a few months post-transplant, with no discernible systematic time-dependence. Therefore, we consider the steady-state solutions to our model (Eqs. 1 and 5). Henceforth, all quantities will be assumed to be those at steady-state. First, we can start from n = 1 and inductively solve for the steady-state form of Eqs. 5 to find

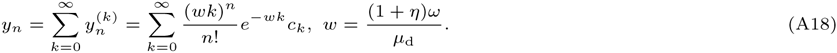

Before this solution can be effectively used in Eq. A10, an explicit expression for the steady-state progenitor clone size distribution *c_k_* is needed.

The total steady-state progenitor population is given by the solution to *αC* + [*r*(*N*_p_) − *μ*]*N*p = 0. The population-limited growth rate is given by Eq. 3 and the steady-state progenitor cell population is explicitly

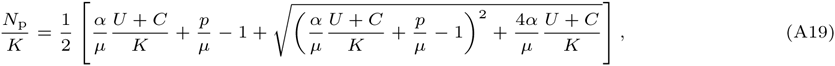

where *μ* = *μ*_p_ + *ηω*. After using Eq. A19 in Eq. 3, we find explicitly

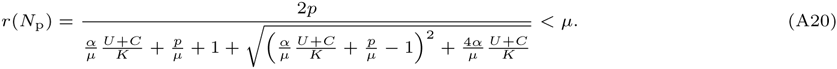

The total differentiated cell population found from 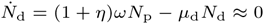 is *N*_d_ = (1 + *η*)*ωN*_p_/*μ*_d_ ≡ *wN*p. Upon using these expressions in the steady-state limit of Eq. 1, we obtain

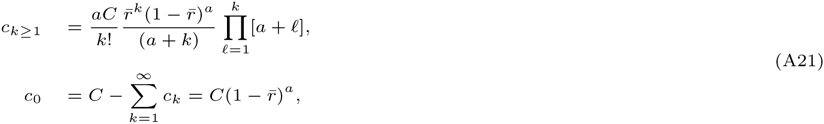

where

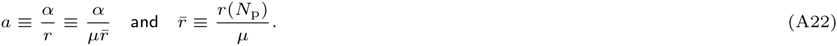

From these results, the total number of clones in each compartment can be explicitly found:

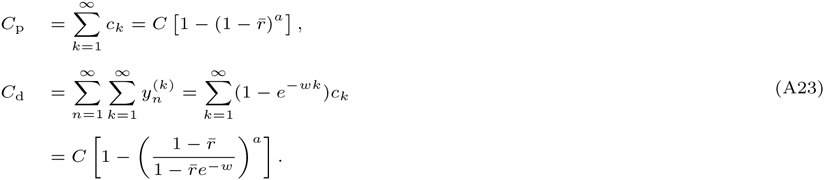

For the total expected number of clones observed in the sample,

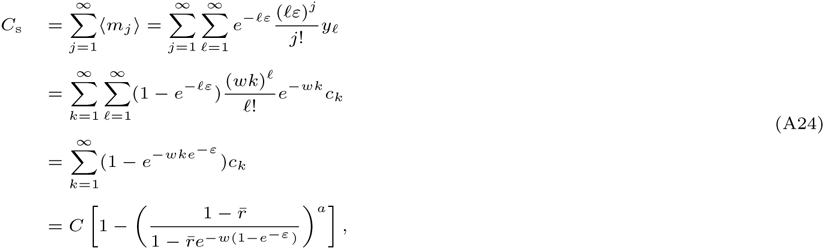

where we have explicitly used Eq. A18 for *y_ℓ_* and Eq. A21 for *c_k_*. This result can be further reduced in two limits

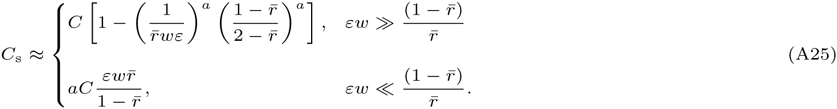

As expected, the total numbers of clones present in each pool follow the progression

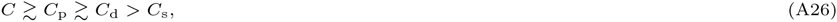

with significant loss of clones due to sampling (*C*_d_ ≫ *C*_s_) only in the second case of Eq. A25 describing sample sizes 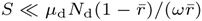. Note that for clone sizes *k* ≫ *a*, *c_k_* in Eq. A21 can be approximated by

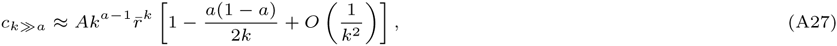

where

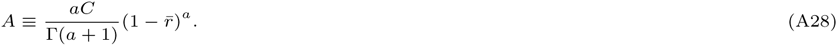

Finally, in steady-state, using Eq. A18 in Eq. A10, we find the cumulative clone size distribution

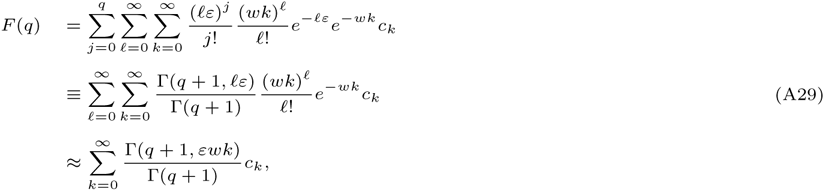

where a steepest descents approximation in *ε* ≪ 1 was used to derive the final approximation. The expected cumulative clone size frequency *Q*(*q*) is obtained by subtracting off the unrepresented clones 〈m_0_〉 ≡ *F*(0) and normalizing by the total expected number of clones *C*_s_ = *F*(*S*) − *F*(0).

A number of numerical procedures can be used to evaluate Q(q) using the final approximation in Eq. A29. For large values of *q*, and small *εw*, the ratio of Γ—functions is near unity for *k* ≲ (*q* + 1)/(*εw*), and quickly decreases to zero for larger *k*. One approach for numerically evaluating *F*(*q*) is to explicitly separate small *q* and small *k* terms in the sum. For small *k*, the exact form of *c_k_* should be used. For larger *k*, the asymptotic form (Eq. A27) can be used and the sum can be approximated as an integral. We find that even for small values *q* ≈ 5, this approximation results in a relative error < 1%. Even the crudest approximation of replacing the sum by the integral

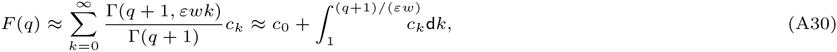

and using the asymptotic form Eq. A27, provides a reasonable estimate of *F*(*q*). This rough approximation also provides an informative analytical expression:

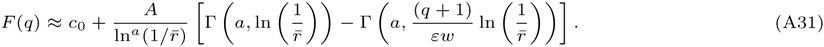

This approximation shows that our distributions depend most strongly on only *a* = *α*/*r* and 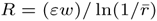 (Eq. 10). Since 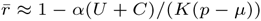 is only very slightly smaller than unity, 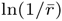 is a small positive number and 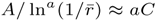. Since 〈*m_q_*〉 = *F*(*q*) − *F*(*q* − 1), an approximate form useful for estimating the clone size distribution is

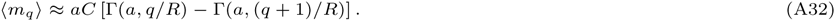

Within physiologically-relevant regimes, our data can be well-fitted to 〈*m_k_*〉 by varying just *a* and *R*. The other physiological parameters, *K*, *p*, *μ*, *U* + *C*, etc., are then related to each other through the most likely numerical values *a*^*^ and *R*^*^ found from fitting the data.

Consider the total number of active HSC cells, *U* + *C*, and the ratio of the rate of HSC differentiation to the rate of self-renewal of progenitor cells, *α*/*p*. Once the best fit parameters *a*^*^ and *R*^*^ have been estimated from fitting clonal frequency distributions, *U* + *C* and *α*/*p* can be expressed in terms of *K*, *S*, and Δ ≡ *p*/*μ* − 1. Note that *S*, the number of sequencing reads detected in each sample, is an experimentally determined parameter.

To find these relationships, we first assume that *εw*/*R*^*^ = (*S/N*_d_)(*ω*/*μ*_d_)/*R*^*^ = *S*/(*R*^*^*N*_p_) ≪ 1 and *S*/(*R*^*^*K*) ≪ 1. By using the definition of *R*, we find 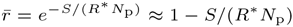. Since 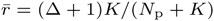 also, these two independent expressions for 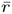 furnish a quadratic equation for *N*_p_ in terms of *R*^*^. After comparing the positive root of this equation to the definition of the steady-state progenitor population *N*_p_ (Eq. A19), we find

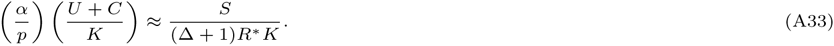

Eq. A33 can be then used to find an expression for *N*_p_ that is independent of *α/p* and *U* + *C*. Using this form of *N*_p_ in the definition *a* = *α/r* = (*α*/*p*) [*N*_p_/*K* + 1], we find an explicit expression for the best-fit value *α*/*p* = *a*^*^*K*/(*N*_p_ + *K*). Further assuming that *S*/(*R*^*^*K*) ≪ Δ, we find

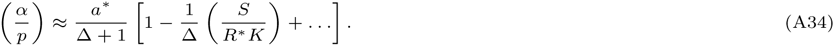

Note that to lowest order, *α*/*p* can be estimated from *a*^*^ and Δ = *p*/*μ* − 1. Finally, substituting (*α*/*p*) ≈ *a*^*^/(Δ + 1) from solving Eq. A34 into Eq. A33, we find _S_

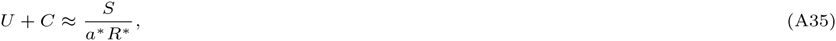

which is independent of *K* and Δ. Note that these parameters can be extracted out of the many parameters in the model because of the limiting values of 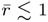 and *εw* ≪ 1. Our model allows one to make predictions on the number of expected clones in each pool, *C*_p_, *C*_d_, and the measured *C*_s_ (Eqs. A23-A24), as well as expected clone size distributions (Eq. A32) as functions of sampling fraction *ε*, turnover rate *w*, effective differentiation rate *a*, and effective growth rate 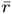. However, from the functional forms of *C*_p_, *C*_d_, *C*_s_, and because 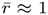, the numerical determination of the number of clones in each pool is highly sensitive to high values of *R* and low values of *a*.

Expected clone size distributions from stochastic clones sizes:

Here, we explicitly show how the neutral assumption (identical transition rates and fitness for all clones) of our populations allows mean-field equations for the expected clone size distribution to be derived from considerations of the stochastic dynamics of an individual clone. Analysis of individual clones is more natural in settings where each clone can be easily isolated and imaged, such as in epidermal systems and geometries [42, 54, 55]. An important feature of our neutral model is that the steady-state clone size distribution depends on only the value of the effective growth rate at steady-state and not on the specific form of the regulation. In other words, the *relative* sizes of neutral clones are independent of the growth law common to all clones. Therefore, we first consider the corresponding birth-death process of a single isolated clone in the presence of constant immigration occurring at rate *α*. The master equation for the probability *p_k_* (*t*) of a single clone containing *k* progenitor cells is

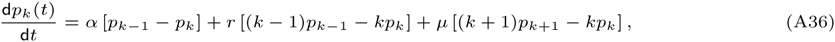

where in our application, *μ* ≡ *μ*_p_ + *ηω*. If the growth rate *r* is assumed constant and independent of *k*, an analytic expression expressed in terms of the corresponding generating function 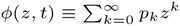 [45, 56]:

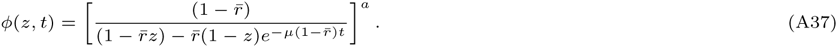

We now identify *c_k_*(*t*) with *C* times the probability that any independent clone is of size *k*. Thus,

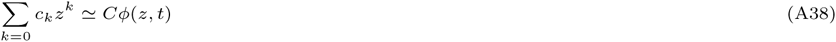

and the variability in clone sizes arises from the variability of the times of differentiation of HSC cells to create progenitor cells of different lineages. In the *t* → ∞ steady-state limit, we find

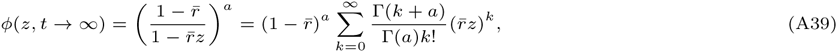

in which 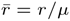. Thus, the single stochastic clone construction of the expected clone size distribution yields

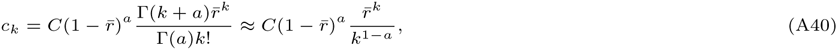

which matches the result in Eq. A27. This derivation explicitly shows that the exponentially distributed initial differentiation times sets the progenitor cell clone size distribution *c_k_*. This distribution is preserved even in the mean-field setting of the hodograph-transformed model described by Eq. 1 and is independent of the specific form chosen for the growth law *r*.

HSC self-renewal:

Rather than assuming that HSCs differentiate only asymmetrically, leaving each unique HSC clone unchanged, we now consider the effects of symmetric HSC replication on the measured clone size distribution. We also assume a separate HSC niche with a corresponding carrying capacity *K*_s_. If we denote *x_k_* as the number of clones in the stem cell niche that is represented by exactly *k* stem cells,

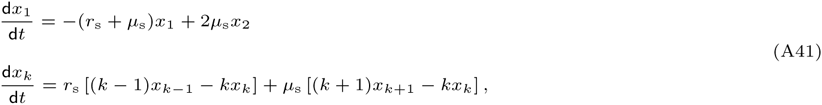

where the effective growth rate *r*_s_ is defined by the carrying capacity *K*_s_, and the total number of stem cells *N*_s_, labeled and unlabeled, in the stem cell compartment:

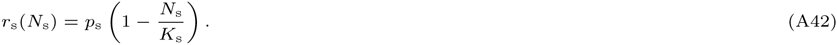

Here, we have used logistic growth for mathematical convenience and to simply illustrate the insensitivity of the final clone size distribution to the model of HSC differentiation. The total stem cell population is defined as 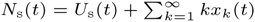. Upon summing Eqs. A41 and the equation for unlabeled cells, 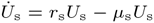, we find that the total population decouples and obeys

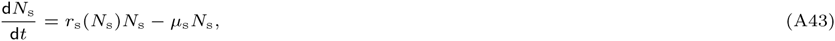

which can be solved exactly, allowing one to find *r*_s_ explicitly as a function of time. Eqs. A41 can then be solved numerically to find the stem cell clone frequencies in the stem cell compartment. To simplify the calculations and find a tractable solution, we will set *μ*_s_ = 0 and define a new time variable d*τ* = *r*_s_(*t*)d*t*. Equations A41 for *x_k_*(*τ*) now have constant coefficients and can be solved by using the initial conditions *x*_1_(*τ* = 0) = *C*_s_, *x_k_* _> 1_ (0) = 0, and the Laplace transforms,

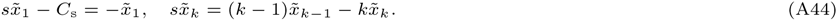

The solution

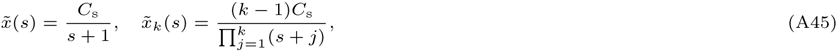

can be inverted to yield

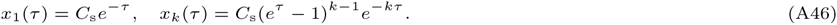

To transform back to *x_k_*(*t*), we need to invert

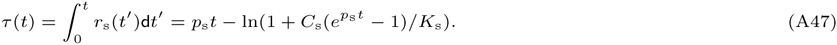

In the steady-state limit, *t* → ∞ corresponds to 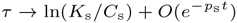. In this limit, Eq. A46 yields

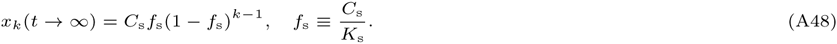

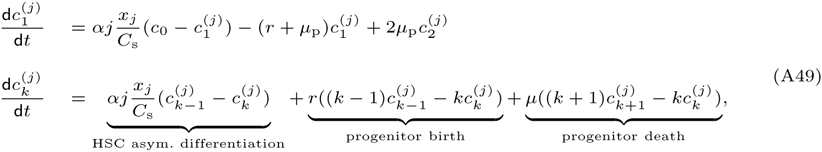

where 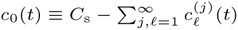 is the total number of clones that do not appear in the progenitor cell pool at time *t*. Upon summing all the above equations to find the zeroth and first moments of 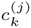, we find

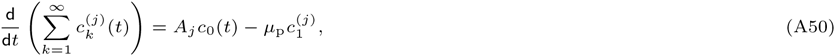

and

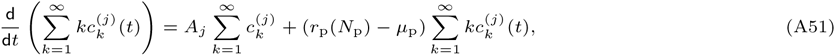

where *A_j_*(*t*) = *αjx_j_*(*t*)/*C*_s_. By further adding 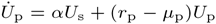 to Eq. A51, we find

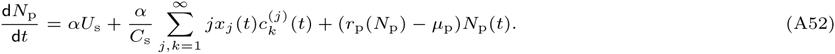

If we assume steady-state in both the stem cell and progenitor cell populations, *A_j_* = *jαx_j_* (∞)/*C*_s_ = *jαf*_s_ (1 − *f*_s_)*^j^*^−1^ and Eq. A50 yields

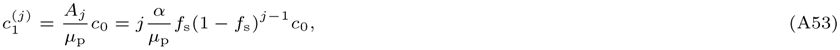

which, when used in the steady-state limit of Eq. A51 yields

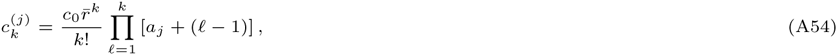

where

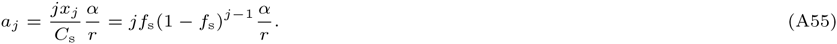

The coefficient *c*_0_ can now be self-consistently calculated by noting that 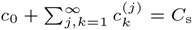. Upon double-summing Eq. A54, we find *c*_0_ = *C*_s_/*Ƶ*, where

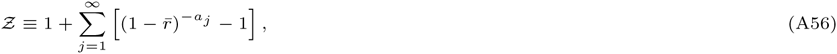

and

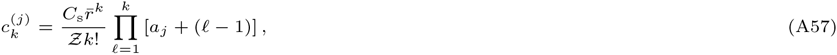

The clone numbers in the progenitor pool are thus 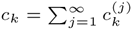. Note that when the initial transplantation fills the entire stem cell niche, 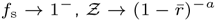, and the only term in Eq. A54 that survives is *j* = 1, leading to our previous result as expected. For the general product in Eq. A54, we can approximate

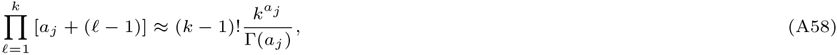

when *a_j_* ≪ ln *k*. From Eq. A55, we know that *a_j_* is strictly bounded above by *α*/*r* and is typically ≲ 0.5*α*/*r* for *f*_s_ < 0.5. Since we expect *α*/*r* < 1, the approximation in Eq. A58 is valid for essentially all values of *k* ≳ 2. In order to compute *c_k_*, we perform the sum

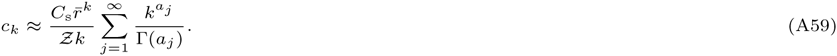

By Taylor-expanding in small *a_j_* first, we can further approximate the sum as

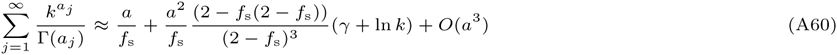

where *a* ≡ *α*/*r* and *γ* ≈ 0.5772 is Euler’s constant. In order to find an explicit expression for *Ƶ*, note that 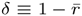 is typically very small, on the order of 1/*K*_p_. Therefore 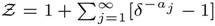 can be approximated using 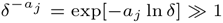 and Laplace’s method on the sum. The dominant term in the sum arises for *j*^*^ ≈ −1/ ln(1 − *f*_s_). Approximating the sum by an integral over a Gaussian centered about *j*^*^, we find

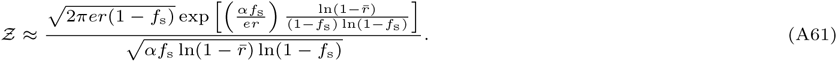

By using this approximation for *Ƶ* and Eq. A60 in Eq. A59, we can find the leading behavior of *c_k_* in the small *f*_s_ and *a* ln *k* ≪ 1 limits,

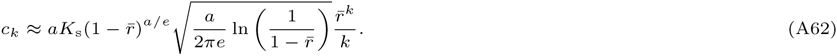

This approximation is not good for small *a* since it suppresses the largeness of the asymptotic parameter 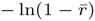. Instead, for small 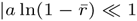, we Taylor expand exp[−*a_j_* ln *δ*] ≈ 1 − *a_j_* ln *δ* to find

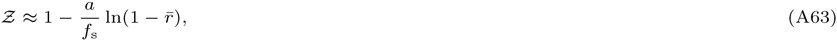

and

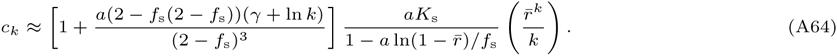

Thus, in the limit where the transplanted number of clones *C*_s_ ≪ *K*_s_ is much less than the HSC carrying capacity, the marked HSCs greatly expand before reaching *K*_s_; however, the resulting clone size distribution *c_k_* remains qualitatively unchanged.

Maximum Likelihood Estimation:

We start with the counts of unique sequencing reads on the macaque genome. *i.e.* number of times the read was sequenced. We refer to each unique read as a “clone.” Since sequencing of each end of a unique viral sequence is performed independently, we treat the two data sets as independent measurements at each time. The reads are then pooled according to which end of a read was sequenced. For more details of the sequencing and filtering of the reads, see Kim *et al.* [14].

Assume a sequencing run from a particular animal, at a particular time and at one of the sequencing ends, yields *n* unique clones with {*q*_1_, …, *q_i_*,…,*q_n_*} read counts. We calculate the likelihood of observing this data within our model given a particular set of parameter values. Our mathematical model contains three independent parameter combinations:

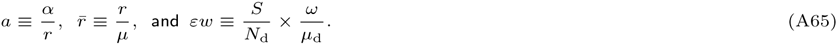

Since the distinguishable clones are otherwise physiologically identical, we associate the distribution of sizes of any particular clone with the expected value of clone size distribution:

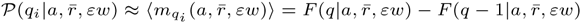. The likelihood of the parameters given the detected clone sizes {*q*_1_, …, *q_n_*} is then given by:

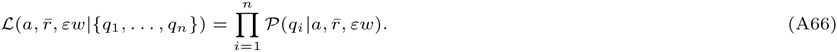

The most likely parameters are then estimated by numerically maximizing the likelihood over the parameters. However, as shown previously, the distribution of clone sizes depends most strongly on only *a* and *R* given by Eqs. 10.

The figures below show normalised and rescaled clone size distributions extracted from granulocyte or peripheral blood mononuclear cell (PBMC) subpopulations of blood from all animals in the original study. The MLE values of *a*^*^ and *R*^*^ all fall within regimes such that *U* + *C* ~ 10^3^ − 10^4^. The fluctuations in *U* + *C* are predominantly due to changes in the fraction *f* at different time points. Such fluctuations are the result of internal dynamics not considered in our model and do not exhibit any discernible trend.

**Fig. A1.**
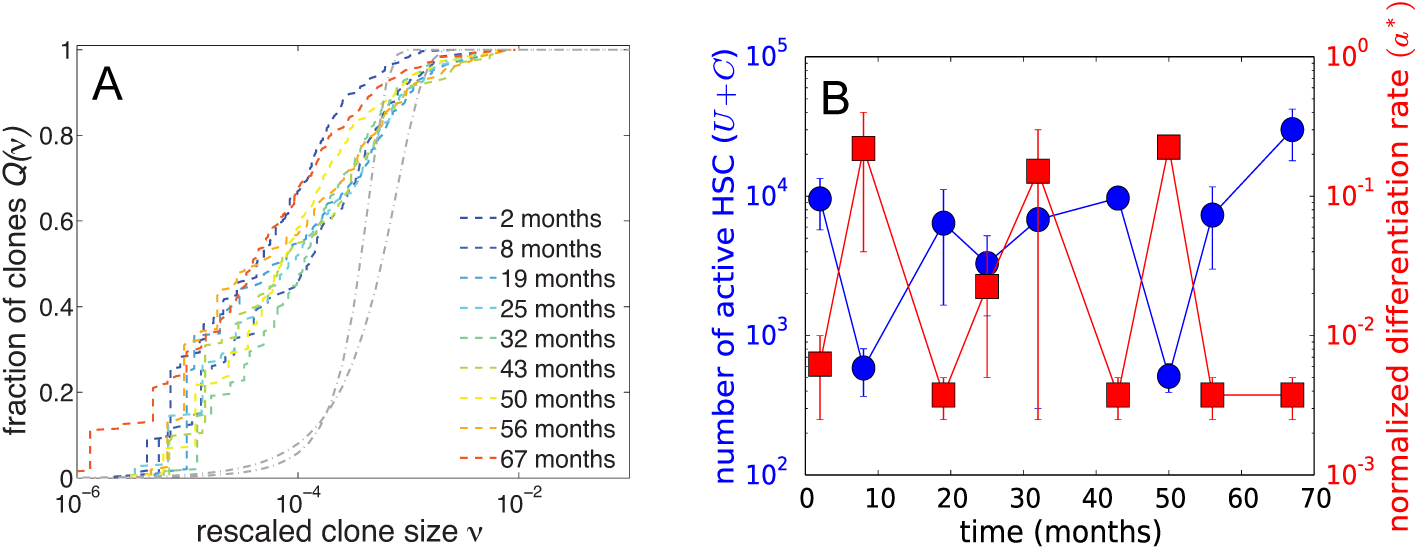
(A) Rescaled clone size distributions from granulocytes in animal RQ5427. The stationarity in this subpopulation is evident. (B) The MLE fits for *U* + *C* (blue circles) and *a*^*^ (red squares).

**Fig. A2.**
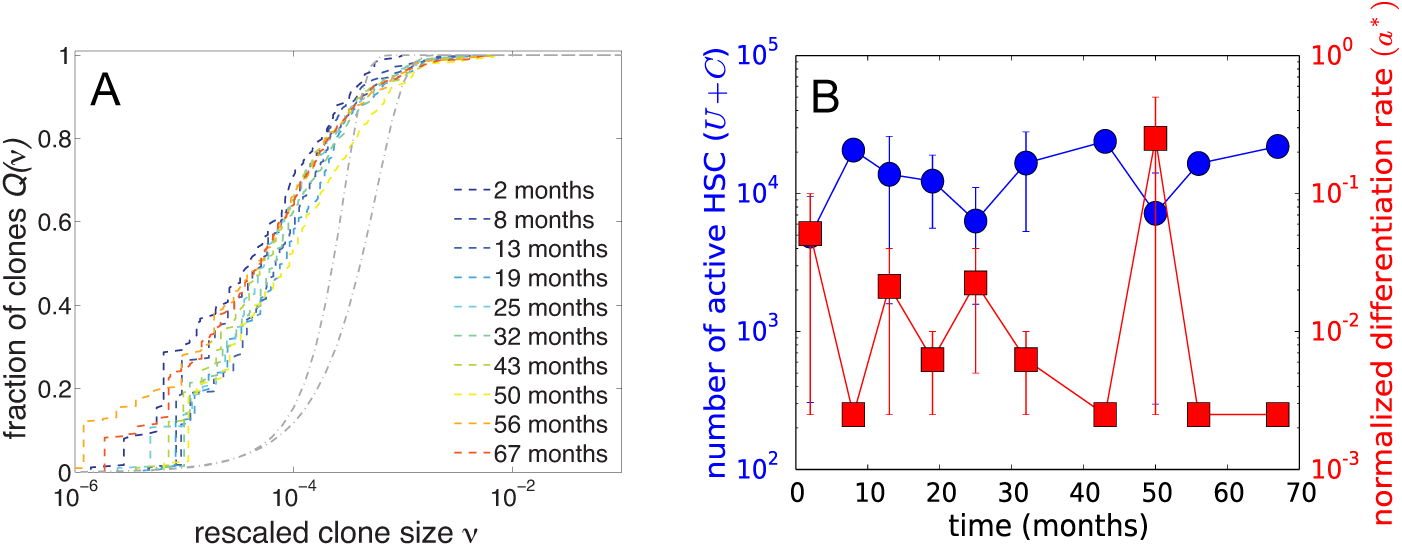
(A) Rescaled PBMC clone size distributions from animal RQ5427. (B) MLE fits for *U* + *C* and *a*^*^.

**Fig. A3.**
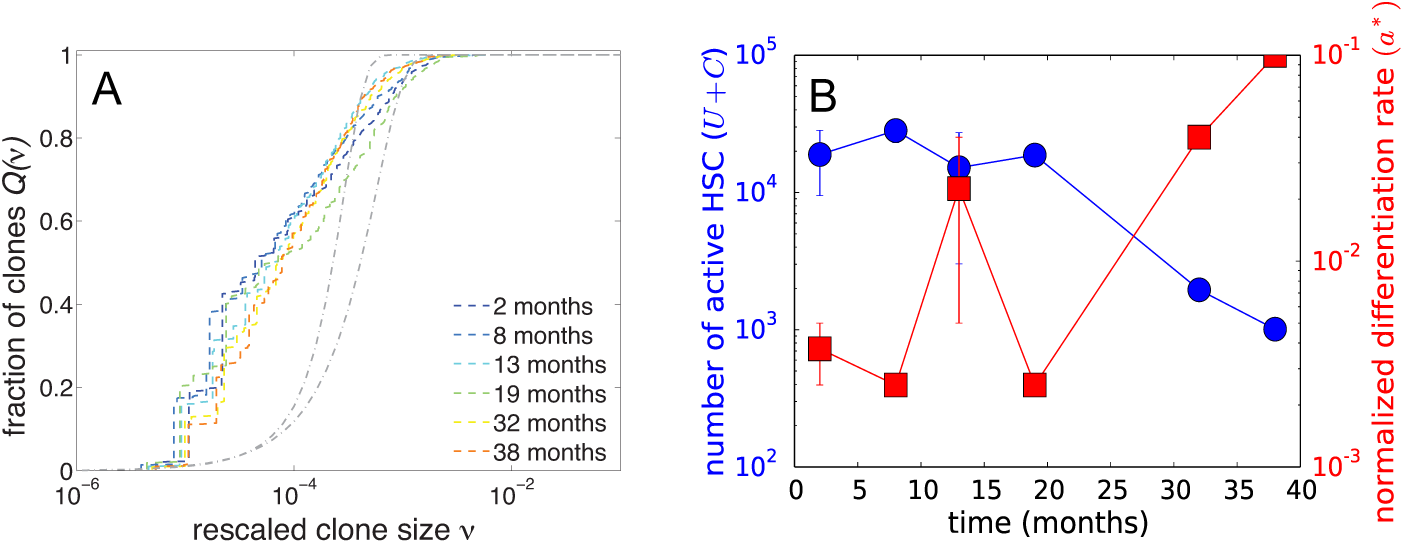
(A) Rescaled granulocyte clone size distributions from animal RQ3570. (B) MLE fits for *U* + *C* and *a*^*^.

**Fig. A4.**
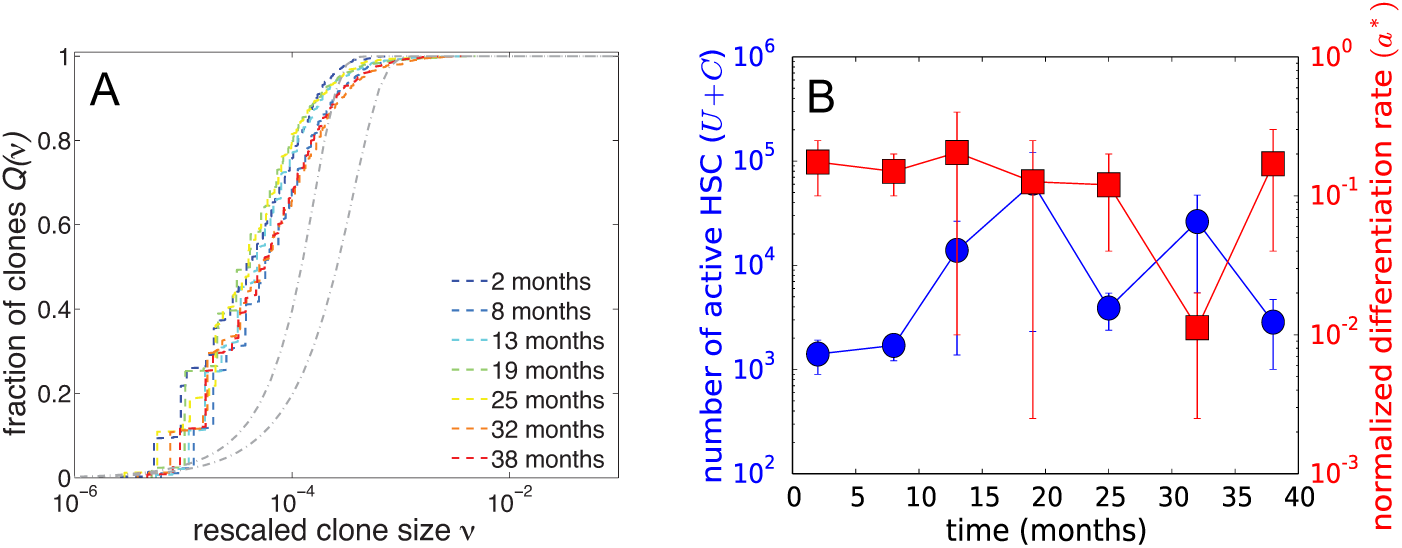
(A) Rescaled PBMC clone size distributions from animal RQ3570. (B) MLE fits for *U* + *C* and *a*^*^.

**Fig. A5.**
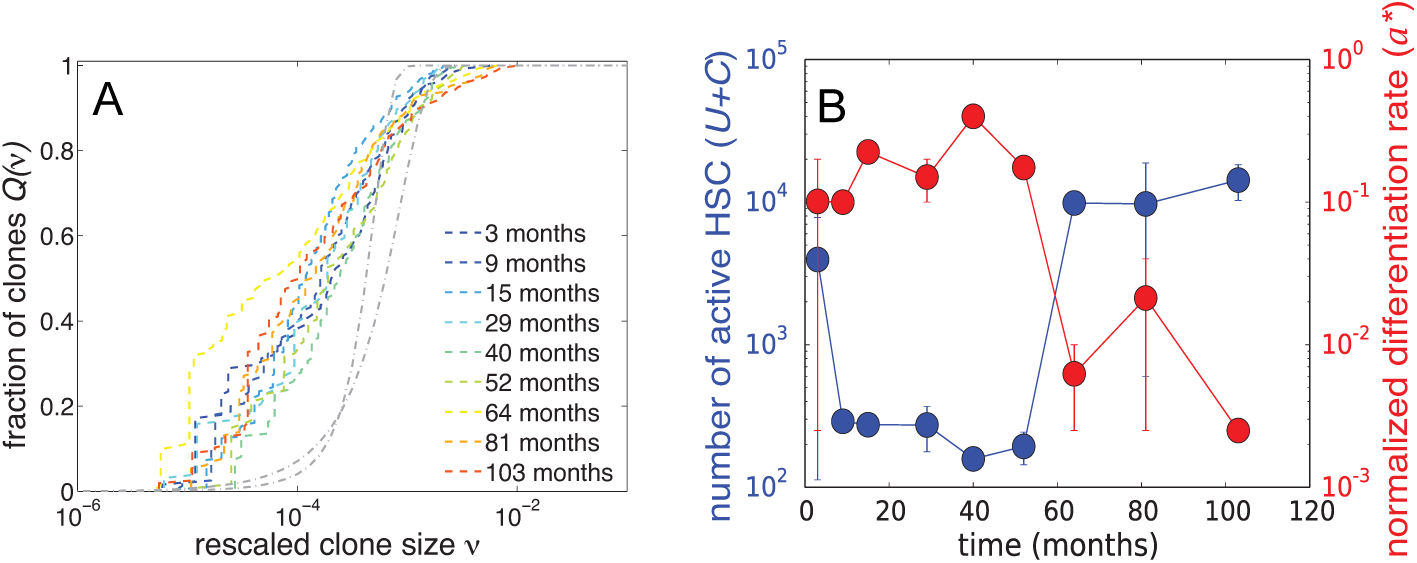
(A) Rescaled clone size distributions from granulocytes in animal 2RC003. The stationarity in this subpopulation is evident. (B) The MLE fits for *U* + *C* (blue circles) and *a*^*^ (red squares).

**Fig. A6.**
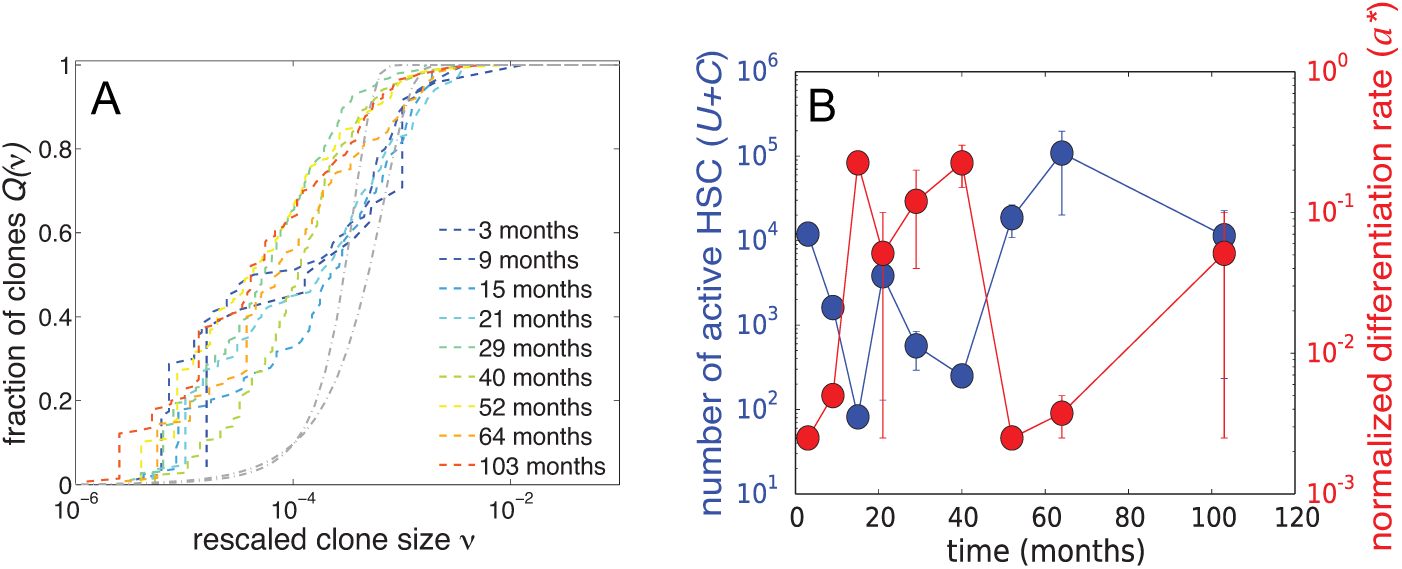
(A) Rescaled PBMC clone size distributions from animal 2RC003. (B) MLE fits for *U* + *C* and *a*^*^.

**Fig. A7.**
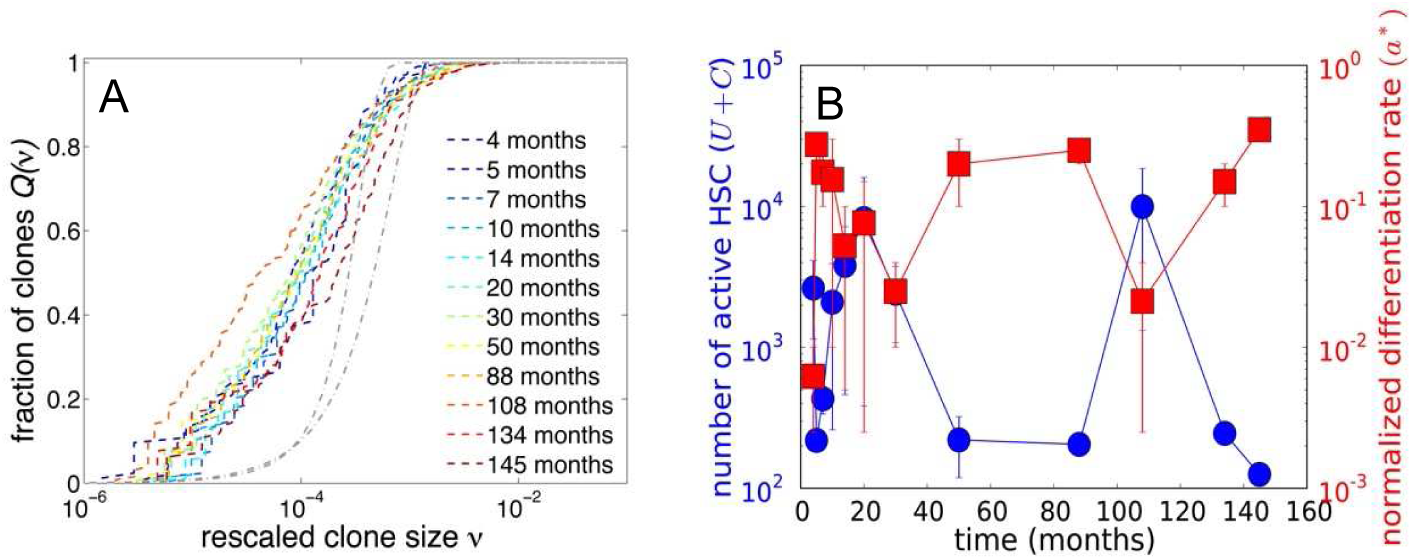
For animal 95E132 clone sizes were measured only for within the total peripheral blood cells (PBC) population. (A) Rescaled clone size distributions from animal 95E132. (B) MLE fits for *U* + *C* and *a*^*^ from the combined PBC clone size distributions.

**Fig. A8.**
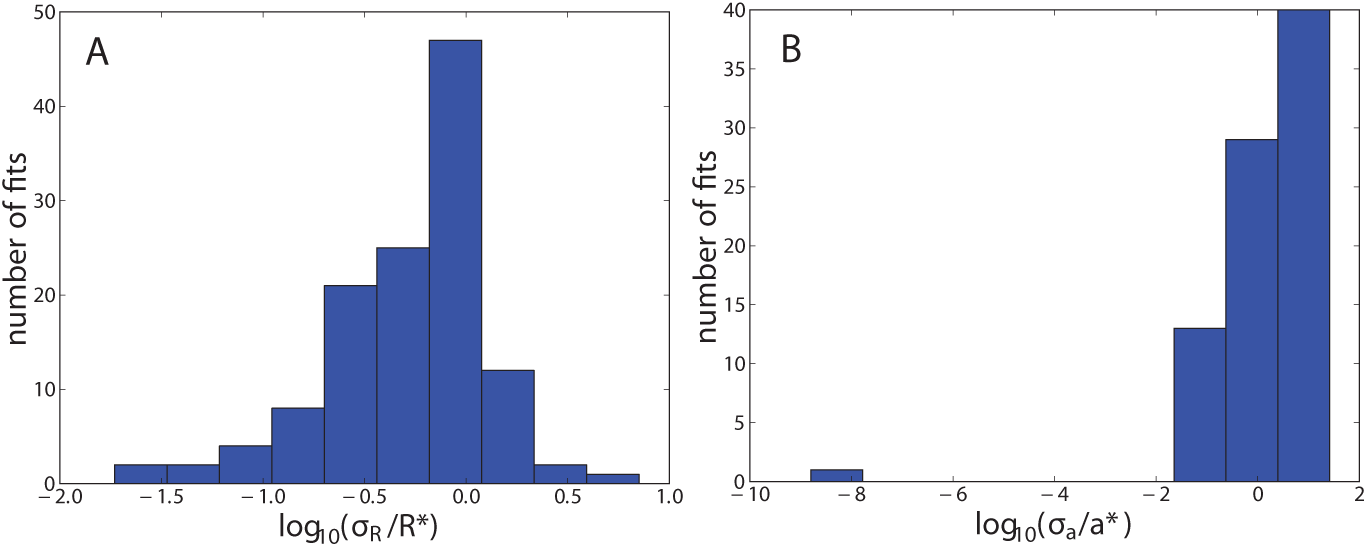
(A) Distribution of log(*σ_R_*/*R*^*^) values across all data sets. (B) Distribution of log(σ*_a_*/*a*^*^) values.

For completeness, we also calculate a rough goodness-of-fit metric. We do this by calculating the “diagonal” curvatures of the likelihood function 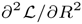 and 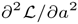 evaluated at the maximum (*R*^*^, *a*^*^). Upon defining

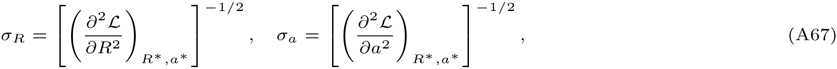

a goodness-of-fit can be measured through the distribution of the values of the Fano factors *σ_a_*/*a*^*^ and *σ_R_*/*R*^*^ obtained by fitting each clone size distribution at each time point. The distributions of the logarithm of *σ_a_*/*a*^*^ and *σ_R_*/*R*^*^ (sampled from fitting at all times points for all animals) are plotted below. We see that the fitting for *R* at most time points is reasonably good, but that some of the fits, particularly for the small values of *a*^*^, are not particularly well-conditioned.

## References

1. Enver, T., Heyworth, C.M., Dexter, T.M.: Do stem cells play dice? Blood 92(2), 348–352 (1998)

2. Hoang, T.: The origin of hematopoietic cell type diversity. Oncogene 23(43), 7188–7198 (2004)

3. Muller-Sieburg, C.E., Cho, R.H., Thoman, M., Adkins, B., Sieburg, H.B.: Deterministic regulation of hematopoietic stem cell self-renewal and differentiation. Blood 100, 1302–1309 (2002)

4. Copley, M.R., Beer, P.A., Eaves, C.J.: Hematopoietic stem cell heterogeneity takes center stage. Cell stem cell 10(6), 690–697 (2012)

5. Muller-Sieburg, C.E., Sieburg, H.B., Bernitz, J.M., Cattarossi, G.: Stem cell heterogeneity: implications for aging and regenerative medicine. Blood 119(17), 3900–3907 (2012)

6. Lu, R., Neff, N.F., Quake, S.R., Weissman, I.L.: Tracking single hematopoietic stem cells in vivo using high-throughput sequencing in conjunction with viral genetic barcoding. Nature Biotechnology 29(10), 928–933 (2011)

7. Huang, S.: Non-genetic heterogeneity of cells in development: more than just noise. Development 136, 3853–3862 (2009)

8. Osafune, K., Caron, L., Borowiak, M., Martinez, R.J., Fitz-Gerald, C.S., et al.: Marked differences in differentiation potential among human embryonic stem cell lines. Nature Biotechnology 26, 313–315 (2008)

9. Pang, W.W., Price, E.A., Sahoo, D., Beerman, I., Maloney, W.J., Rossi, D.J., Schrier, S.L., Weissman, I.L.: Human bone marrow hematopoietic stem cells are increased in frequency and myeloid-biased with age. Proc. Natl. Acad. Sci. USA 108, 20012–20017 (2011)

10. Harrison, D.E., Astle, C.M., Lerner, C.: Number and continuous proliferative pattern of transplanted primitive immunohematopoietic stem cells. Proc. Natl. Acad. Sci. USA 85(3), 822–826 (1988)

11. Verovskaya, E., Broekhuis, M.J.C., Zwart, E., Ritsema, M., van Os, R., de Haan, G., Bystrykh, L.V.: Heterogeneity of young and aged murine hematopoietic stem cells revealed by quantitative clonal analysis using cellular barcoding. Blood 122(4), 523–532 (2013)

12. Takizawa, H., Regoes, R.R., Boddupalli, C.S., Bonhoeffer, S., Manz, M.G.: Dynamic variation in cycling of hematopoietic stem cells in steady state and inflammation. J. Exp. Med. 208(2), 273–284 (2011)

13. Gerrits, A., Dykstra, B., Kalmykowa, O.J., Klauke, K., Verovskaya, E., Broekhuis, M.J.C., de Haan, D., Bystrykh, L.V.: Cellular barcoding tool for clonal analysis in the hematopoietic system. Blood 115(13), 2610–2618 (2010)

14. Kim, S., Kim, N., Presson, A.P., An, D.S., Mao, S.H., Bonifacino, A.C., Donahue, R.E., Chow, S.A., Chen, I.S.Y.: High-Throughput, Sensitive Quantification of Repopulating Hematopoietic Stem Cell Clones. Journal of Virology 84(22), 11771–11780 (2010)

15. Biffi, A., Montini, E., Lorioli, L., Cesani, M., Fumagalli, F., et al.: Lentiviral hematopoietic stem cell gene therapy benefits metachromatic leukodystrophy. Science 341(6148), 1233158 (2013)

16. Aiuti, A., Biasco, L., Scaramuzza, S., Ferrua, F., Cicalese, M.P., et al.: Lentiviral hematopoietic stem cell gene therapy in patients with Wiskott-Aldrich syndrome. Science 341(6148), 1233151 (2013)

17. Cavazzana-Calvo, M., Payen, E., Negre, O., Wang, G., Hehir, K., et al.: Transfusion independence and HMGA2 activation after gene therapy of human *β*-thalassaemia. Nature 467(7313), 318–322 (2010)

18. Cartier, N., Hacein-Bey-Abina, S., Bartholomae, C.C., Veres, G., Schmidt, M., et al.: Hematopoietic stem cell gene therapy with a lentiviral vector in X-linked adrenoleukodystrophy. Science 326(5954), 818–823 (2009)

19. Kim, S., Kim, N., Presson, A.P., Metzger, M.E., Bonifacino, A.C., et al.: Dynamics of HSPC Repopulation in Nonhuman Primates Revealed by a Decade-Long Clonal-Tracking Study. Cell Stem cell 14(4), 473–485 (2014)

20. Sun, J., Ramos, A., Chapman, B., Johnnidis, J.B., Le, L., Ho, Y.-J., Klein, A., Hofmann, O., Camargo, F.D.: Clonal dynamics of native haematopoiesis. Nature 514, 322–327 (2014)

21. Loeffler, M., Roeder, I.: Tissue stem cells: definition, plasticity, heterogeneity, self-organization and models - a conceptual approach. Cells Tissue Organs 171, 8–26 (2002)

22. Bernard, S., Belair, J., Mackey, M.C.: Oscillations in cyclical neutropenia: New evidence based on mathematical modeling. J. Theor. Biol. 223(3), 283–298 (2003)

23. Dingli, D., Pacheco, J.M.: Modeling the architecture and dynamics of hematopoiesis. Wiley Interdisciplinary Reviews. Systems biology and medicine 2(2), 235–244 (2010)

24. Courant, R.: Differential and Integral Calculus, Vol. II. Blackie & Son, London (1936)

25. D’Orsogna, M.R., Lakatos, G., Chou, T.: Stochastic self-assembly of incommensurate clusters. J. Chem. Phys. 136, 084110 (2012)

26. Marciniak-Czochra, A., Stiehl, T., Ho, A.D., Jager, W., Wagner, W.: Modeling of asymmetric cell division in hematopoietic stem cells-regulation of self-renewal is essential for efficient repopulation. Stem Cells and Development 18, 377–385 (2009)

27. Kent, D.G., Li, J., Tanna, H., Fink, J., Kirschner, K., Pask, D.C., et al.: Self-renewal of single mouse hematopoietic stem cells is reduced by JAK2V617F without compromising progenitor cell expansion. PLoS Biology 11, 1001576 (2013)

28. Lander, A.D., Gokoffski, K.K., Wan, F.Y.M., Nie, Q., Calof, A.L.: Cell lineages and the logic of proliferative control. PLoS Biology 7, 1000015 (2009)

29. Hoffmann, M., Chang, H.H., Huang, S., Ingber, D.E., Loeffler, M., Galle, J.: Noise-driven stem cell and progenitor population dynamics. PLoS One 3, 2922 (2008)

30. Roshan, A., Jones, P.H., Greenman, C.D.: Exact, time-independent estimation of clone size distributions in normal and mutated cells. J. Roy. Soc. Interface 11, 20140654 (2014)

31. McHale, P.T., Lander, A.: The Protective Role of Symmetric Stem Cell Division on the Accumulation of Heritable Damage. PLoS Computational Biology 10, 1003802 (2014)

32. Antal, T., Krapivsky, P.L.: Exact solution of a two-type branching process: Clone size distribution in cell division kinetics. Journal of Statistical Mechanics, 07028 (2010)

33. Abkowitz, J.L., Caitlin, S.N., McCallie, M.T., Guttorp, P.: Evidence that the number of hematopoietic stem cells per animal is conserved in mammals. Blood 100(7), 2665–2667 (2002)

34. Morozov, A.Y., Bruinsma, R., Rudnick, J.: Assembly of viruses and the pseudo-law of mass action. J. Chem. Phys. 131, 155101 (2009)

35. Krapivsky, P.L., Ben-Naim, E., Redner, S.: Statistical Physics of Irreversible Processes. Cambridge University Press, Cambridge, UK (2010)

36. Szkely, T. Jr, Burrage, K., Mangel, M., Bonsall, M.B.: Stochastic dynamics of interacting haematopoietic stem cell niche lineages. PLoS Computational Biology 10, 1003794 (2014)

37. Mackay, R.: Unified Hypothesis for the Origin of Aplastic Anemia and Periodic Hematopoiesis. Blood 51(5), 941–956 (1978)

38. Keshet-Edelstein, L.: Mathematical Models in Biology. SIAM, New York, NY (2005)

39. Wolfensohn, S., Lloyd, M.: Handbook of Laboratory Animal Management and Welfare, 3rd Edition. Blackwell Publishing, Oxford (2003)

40. Kimura, M.: Population genetics, molecular evolution, and the neutral theory: Selected papers. In: Takahata, N. (ed.) Population Genetics, Molecular Evolution, and the Neutral Theory: Selected Papers. University of Chicago Press, Chicago, IL (1995)

41. He, J., Zhang, G., Almeida, A.D., Cayoutte, M., Simons, B.D., Harris, W.A.: How variable clones build an invariant retina. Neuron 75, 786–798 (2012)

42. Blanpain, C., Simons, B.D.: Unravelling stem cell dynamics by lineage tracing. Nature Reviews, Molecular and Cell Biology 14, 489–502 (2013)

43. Guillaume, T., Rubenstein, D.B., Symann, M.: Immune Reconstitution and Immunotherapy After Autologous Hematopoietic Stem Cell Transplantation. Blood 92, 1471–1490 (1998)

44. Tzannou, I., Leen, A.M.: Accelerating immune reconstitution after hematopoietic stem cell transplantation. Clinical and Translational Immunology 3, 11 (2014)

45. Allen, L.J.S.: An Introduction to Stochastic Processes with Applications to Biology. Pearson Prentice Hall, Upper Saddle, NJ (2003)

46. Shepherd, B.E., Guttorp, P., Lansdorp, P.M., Abkowitz, J.L.: Estimating human hematopoietic stem cell kinetics using granulocyte telomere lengths. Exp. Hematology 32(11), 1040–1050 (2004)

47. Shepherd, B.E., Kiem, H.-P., Lansdorp, P.M., Dunbar, C.E., Aubert, G., LaRochelle, A., Seggewiss, R., Guttorp, P., Abkowitz, J.L.: Hematopoietic stem-cell behavior in nonhuman primates. Blood 110(6), 1806–1813 (2007)

48. Driessens, G., Beck, B., Caauwe, A., Simons, B.D., Blanpain, C.: Defining the mode of tumour growth by clonal analysis. Nature 488(7412), 527–530 (2012)

49. Sieburg, H.B., Cattarossi, G., Muller-Sieburg, C.E.: Lifespan differences in hematopoietic stem cells are due to imperfect repair and unstable mean-reversion. PLoS Computational Biology 9, 1003006 (2013)

50. Weiss, G.H.: Equations for the age structure of growing populations. The Bulletin of Mathematical Biophysics 30(3), 427–435 (1968)

51. Catlin, S.N., Busque, L., Gale, R.E., Guttorp, P., Abkowitz, J.L.: The replication rate of human hematopoietic stem cells in vivo. Blood 117(17), 4460–4466 (2011)

52. DeBoer, R.J., Mohri, H., Ho, D.D., Perelson, A.S.: Turnover Rates of B Cells, T Cells, and NK Cells in Simian Immunodeficiency Virus-Infected and Uninfected Rhesus Macaques. J. Immunology 170, 2479–2487 (2003)

53. Pillay, J., den Braber, I., Vrieskoop, N., Kwast, L.M., de Boer, R.J., Borghans, J.A.M., Tesselaar, K., Koenderman, L.: In vivo labeling with ^2^H2O reveals a human neutrophil lifespan of 5.4 days. Blood 116, 625–627 (2010)

54. Klein, A.M., Doupé, D.P., Jones, P.H., Simons, B.D.: Mechanism of murine epidermal maintenance: Cell division and the voter model. Phys. Rev. E, 031907 (2008)

55. Klein, A.M., Nikolaidou-Neokosmidou, V., Doupé, D.P., Jones, P.H., Simons, B.D.: Patterning as a signature of human epidermal stem cell regulation. Journal of The Royal Society Interface (2011)

56. Chou, T., Wang, Y.: Fixation times in differentiation and evolution in the presence of bottlenecks, deserts, and oases. J. Theor. Biol. 372, 65–73 (2015)

